# An integrated technology for quantitative wide mutational scanning of human antibody Fab libraries

**DOI:** 10.1101/2024.01.16.575852

**Authors:** Brian M. Petersen, Monica B. Kirby, Karson M. Chrispens, Olivia M. Irvin, Isabell K. Strawn, Cyrus M. Haas, Alexis M. Walker, Zachary T. Baumer, Sophia A. Ulmer, Edgardo Ayala, Emily R. Rhodes, Jenna J. Guthmiller, Paul J. Steiner, Timothy A. Whitehead

## Abstract

Antibodies are engineerable quantities in medicine. Learning antibody molecular recognition would enable the *in silico* design of high affinity binders against nearly any proteinaceous surface. Yet, publicly available experiment antibody sequence-binding datasets may not contain the mutagenic, antigenic, or antibody sequence diversity necessary for deep learning approaches to capture molecular recognition. In part, this is because limited experimental platforms exist for assessing quantitative and simultaneous sequence-function relationships for multiple antibodies. Here we present MAGMA-seq, an integrated technology that combines <u>m</u>ultiple <u>a</u>nti<u>g</u>ens and <u>m</u>ultiple <u>a</u>ntibodies and determines quantitative biophysical parameters using deep <u>seq</u>uencing. We demonstrate MAGMA-seq on two pooled libraries comprising mutants of ten different human antibodies spanning light chain gene usage, CDR H3 length, and antigenic targets. We demonstrate the comprehensive mapping of potential antibody development pathways, sequence-binding relationships for multiple antibodies simultaneously, and identification of paratope sequence determinants for binding recognition for broadly neutralizing antibodies (bnAbs). MAGMA-seq enables rapid and scalable antibody engineering of multiple lead candidates because it can measure binding for mutants of many given parental antibodies in a single experiment.

---

The success of AlphaFold2^1^ for predicting structure from sequence has spurred intense interest in deep learning approaches for protein functional prediction. Arguably the largest open prize in protein biotechnology is learning antibody molecular recognition, as this would enable the *in silico* design of developable, high affinity binders against any antigenic surface. Deep learning has been utilized to advance antibody design approaches for overall structure prediction^2,3^, paratope and epitope identification^4^, affinity maturation^5,6^ and antibody sequence humanization^7^. These examples highlight the promise of deep learning approaches but also their limitations. Put simply, unbiased experimental antibody binding datasets do not exist at the scale required for extant deep learning algorithms to capture antibody molecular recognition^8,9^.

Researchers recently assessed the scale of experimental data required for accurate prediction of antibody binding effects upon mutation^9^. Through simulated data, they found that a training dataset comprising hundreds of thousands of unbiased antibody-antigen binding measurements across thousands of diverse antibody-antigen complexes would be sufficient to learn the effect of mutation on binding energetics. The structure of this data – on the order of a few hundred mutational data points per antibody spread across thousands of antibodies targeting diverse antigenic surfaces - suggests a different paradigm than deep mutational scanning approaches^10^, which assess tens of thousands of mutations for individual proteins. Requirements for this new ‘wide mutational scanning’ paradigm include the ability to (i.) determine quantitative monovalent binding energetics, with measurement uncertainty, for multiple antibodies against different antigens and over a wide dynamic range, (ii.) recapitulate the native pairing of variable heavy and light chains which can be achieved using antigen binding fragments (Fabs), (iii.) track multiple mutations per antibody on either or both chains simultaneously, and (iv.) include internal controls for quality control and validation. This technology could also be deployed immediately for current antibody engineering applications, including the reconstruction of multiple probable antibody development pathways^11^, rapid affinity maturation campaigns for multiple leads simultaneously, fine specificity profiling for antibody paratopes, and antibody repertoire profiling against different immunogens.

Current antibody engineering techniques exist but have not demonstrated the ability to generate the depth of data required for learning antibody molecular recognition. Antibody deep mutational scanning using various display techniques has been demonstrated for different task-specific applications but does not provide quantitative binding information. Deep mutational scanning has been used to determine quantitative changes in binding affinity for protein binders but only for a narrow dynamic range^12,13^. TiteSeq^14^ utilizes yeast surface display and next generation sequencing to ascertain quantitative affinities, but has only been demonstrated for a library from one parental antibody single chain variable fragment (scFv)^15^, which can alter the paratope through the constrained folding of heavy and light chains imposed by an inserted linker^16^. Another high-throughput technique demonstrated for one antibody include high-throughput mammalian display^17^. Additional demonstrations^18,19^ exist that have evaluated multiple antibodies and antigens simultaneously but are not high-throughput.

We introduce **MAGMA-seq**, a technology that combines **<u>m</u>**ultiple **<u>a</u>**nti**<u>g</u>**ens and **<u>m</u>**ultiple **<u>a</u>**ntibodies and determines quantitative biophysical parameters using deep **<u>seq</u>**uencing to enable wide mutational scanning of antibody Fab libraries. We demonstrate the ability of MAGMA-seq to quantitatively measure binding affinities, with associated confidence intervals, for multiple antibody libraries. We validated the results of MAGMA-seq with isogenic antibody variant titrations (i.e. labeling isogenic yeast displaying Fabs at various concentrations of antigen and fitting fluorescence measurements to a binding isotherm to extract K_D_). We further demonstrate the utility of MAGMA-seq on a mixed pool of antibody libraries with two distinct antigens, SARS-CoV-2 spike (S1) and influenza hemagglutinin (HA), and recovered the sequence-binding profiles for six antibodies across four distinct protein surfaces. MAGMA-seq facilitates the engineering of antibodies for different applications in parallel: we demonstrate the mapping of potential antibody development pathways, antibody responses to multiple epitopes simultaneously, and identification of paratope sequence determinants for binding recognition for broadly neutralizing antibodies (bnAbs). MAGMA-seq enables rapid and scalable antibody engineering.

## Results

The protocol for MAGMA-seq (**Figure 1a**) starts by generating mutagenic libraries for all antibodies of interest in a Fab format. Fab libraries are subcloned into yeast display vectors each containing a 20 nt molecular barcode; the Fab variant and barcode are paired by sequencing. The library is transformed into yeast, and yeast is grown and induced to surface display the Fabs. The yeast library is sorted at multiple labeling concentrations of antigen(s) by collecting a fixed percentage of yeast cells. After sorting, the collected yeast plasmids are extracted, and the barcode region is sequenced using short-read sequencing. The sequenced data and sorting parameters are then input into a novel computational maximum likelihood estimation (MLE) pipeline to infer most likely biophysical parameters, and associated confidence intervals, for each antibody variant.

**Fig. 1.**
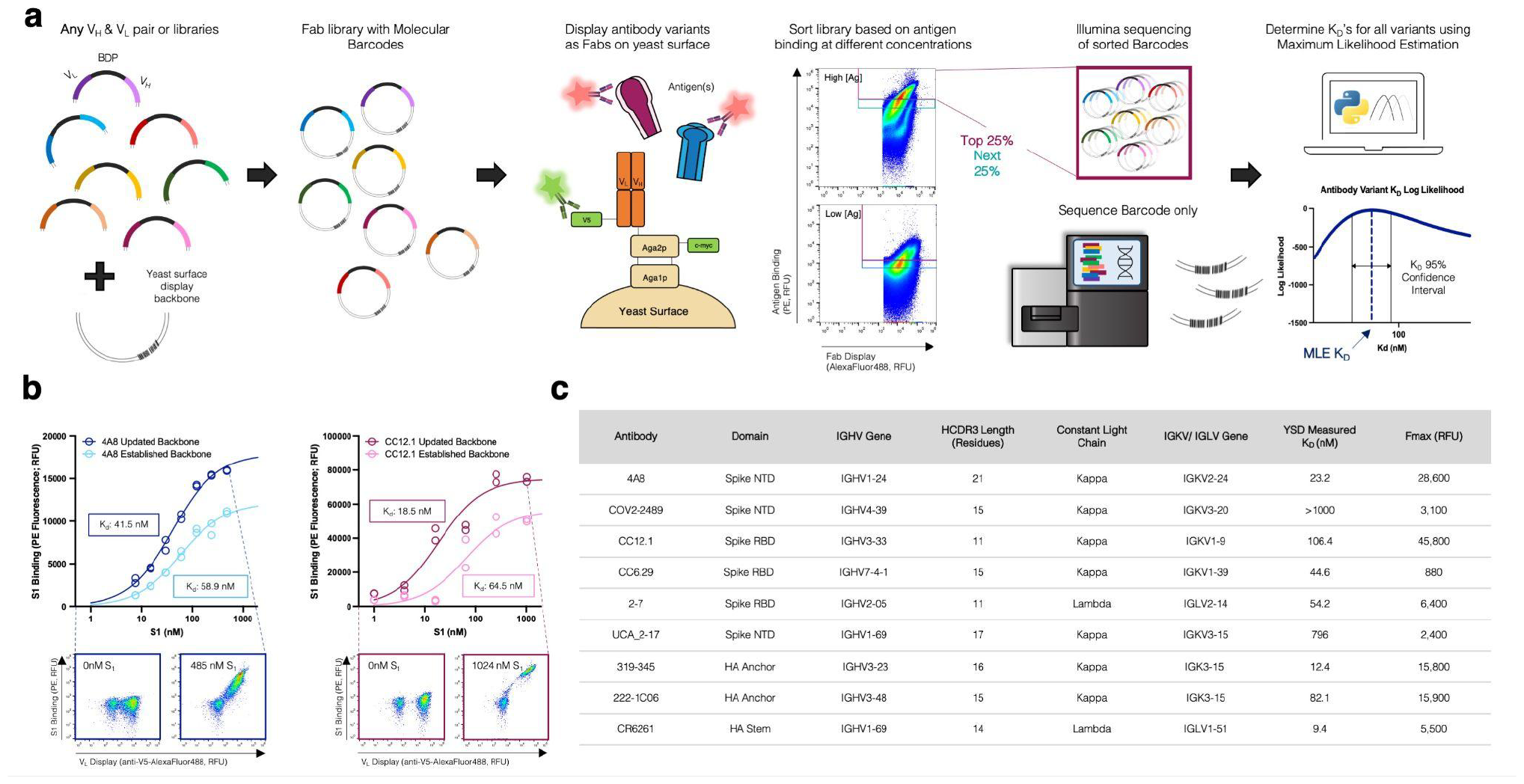
MAGMA-seq is an integrated technology for antibody wide mutational scanning. (a) Protocol schematic. (b) Yeast surface titrations of 4A8 and CC12.1 Fabs against Fc-conjugated S_1_ in the established (light) and updated (dark) yeast surface display vectors. Cytograms from indicated data points are shown for updated yeast surface backbones. Inset describes experimentally determined K_D_ values (n=2). (c) Antibodies assessed using updated yeast surface display vectors. Abbreviations: RBD – Receptor Binding Domain Wuhan Hu-1; NTD – N Terminal Domain Wuhan Hu-1; NA– influenza neuraminidase N2 A/Brisbane/10/2007; HA – influenza hemagglutinin A/Brisbane/02/2018 H1.

There have been several yeast display Fab plasmids described^20–26^; ours most closely relates to a Golden Gate compatible plasmid from Rosowski et al.^20^ Common to many plasmids, including Rosowski et al.^20^, is the light chain and heavy chain (V_H_ and CH1) expressed using a Gal1/Gal10 galactose-inducible bidirectional promoter (BDP). We use Golden Gate^27^ to assemble small shuttle vectors containing the V_H_, V_L_, and BDP, as well as regions of homology to the CH1 and light chain sequence. After mutagenesis, the Fab yeast surface display library is generated by Gibson assembly^28^ using the regions of homology on the shuttle vector and empty yeast surface display vector containing the barcode. Beyond these innovations, we made several useful changes to the Rosowski plasmid (**Extended Data Figure 1**), including (i.) constructing plasmids for both kappa and lambda light chains; (ii.) encoding a V5 C-terminal epitope tag on the light chain to assess light chain expression; and (iii.) making a conservative coding mutant in CH1 and several silent mutations on the yeast vector for compatibility with short-read sequencing.

To test whether our updated plasmids interfered with Fab binding, we performed yeast surface titrations of SARS-CoV-2 antibodies 4A8^29^ and CC12.1^30^ against Wuhan Hu-1 S1 in the established and updated yeast surface display vectors (**Figure 1b**) and fit the mean fluorescence data (F) to a saturable binding isotherm:

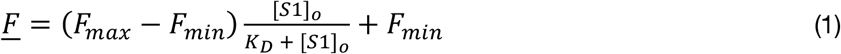

Here F_max_ is the maximum average cell fluorescence at binding saturation, [S1]o is the ligand concentration, F_min_ is the cell autofluorescence, and K_D_ is the monovalent binding dissociation constant. The confidence intervals for K_D_ overlapped for both antibodies (**Fig 1b**), suggesting that the combined changes were not deleterious for binding. For further validation, we performed additional yeast surface titrations with a representative set of antibodies encompassing diverse Complementarity-determining region (CDR) H3 lengths (lengths 11-23), immunoglobulin heavy chain variable region (IGHV) gene families, and either lambda or kappa light chains (**Figure 1c**; **Extended Data Figure 2**). In all cases, interpretable binding isotherms were observed. Thus, our yeast display plasmids can measure binding for a range of human Fabs.

**Fig. 2.**
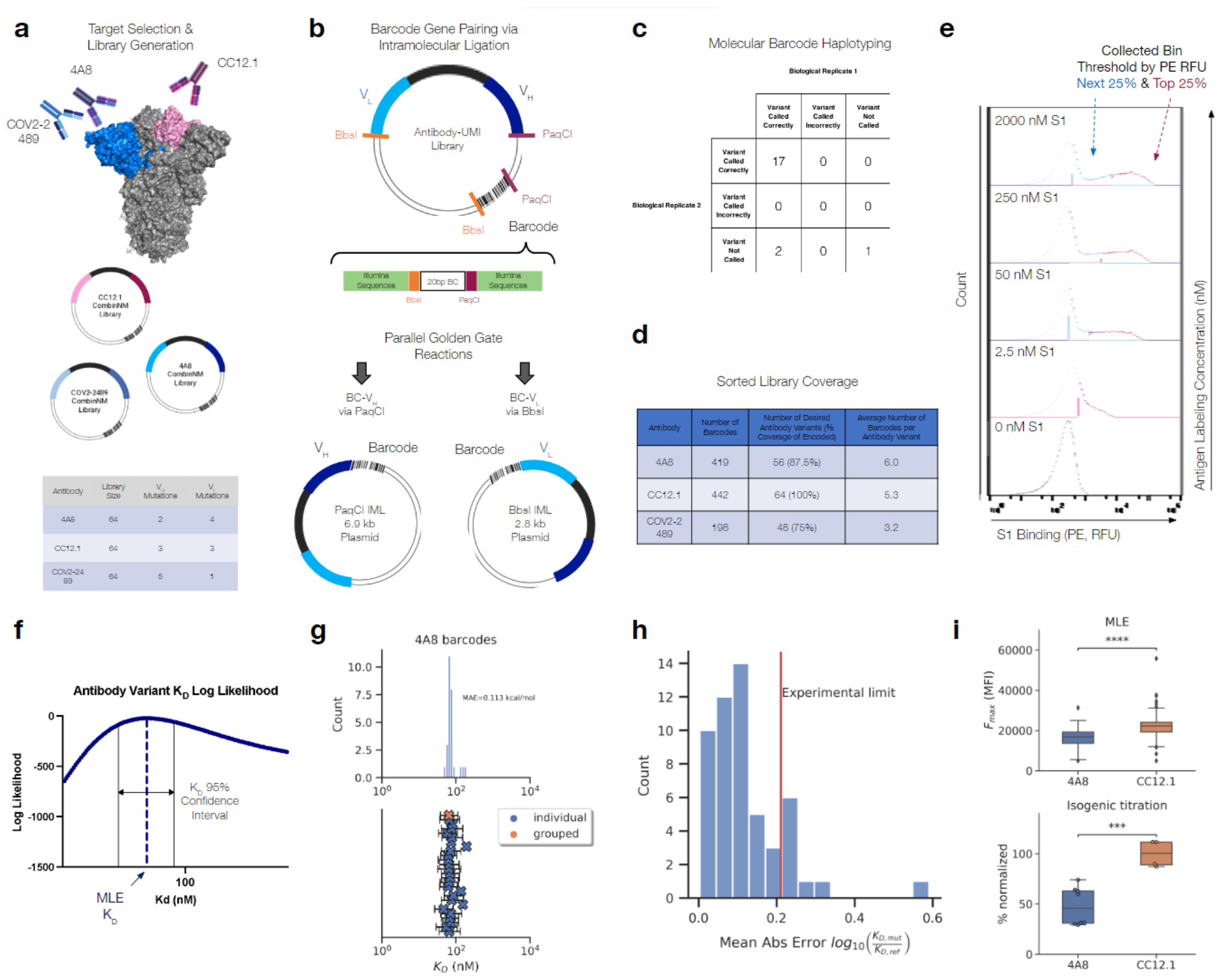
Validation of barcode pairing and parameter estimation for MAGMA-seq. (a) Mutagenic library contains 192 variants of 4A8, COV2-2489 (NTD targeting), and CC12.1 (RBD targeting) Fabs (b) Molecular barcode in yeast display plasmid backbone allows for barcode pairing by intramolecular ligation followed by short-read sequencing (c) Barcode pairing method achieves correct variant calls confirmed by ONT sequencing (d) Barcode and variant coverage of haplotyped libraries (e) Examples of gating thresholds for FACS sorting of library for 4/10 of the sampled antigen concentrations. Top 25% bin shown in pink and next 25% bin shown in blue. (f) MLE quantifies K_D_ uncertainty via confidence interval calculation. (g) MLE K_D_ estimates for all barcodes haplotyped as 4A8 WT (top) with 95% confidence intervals for each barcode (blue X) and grouped barcodes (orange X) (bottom). (h) Mean absolute error for MLE K_D_ estimates for counts collapsed by variant versus isogenic titration values (4A8 only) (i) Maximum mean fluorescence values (F_max_) for 4A8 and CC12.1 antibodies calculated via MLE in absolute terms (top; 4A8: n=70, CC12.1: n=83) and isogenic titration as a percentage normalized by the CC12.1 average (bottom; 4A8: n=8, CC12.1: n=4). P-values calculated by Welch’s t-test (***: 1e-4 < p <= 1e-3, ****: p <= 1e-4).

To demonstrate the capability of MAGMA-seq to track potential development trajectories of multiple antibodies simultaneously, we selected three anti-S1 antibodies^29–31^ that target Wuhan Hu-1 S1 at two distinct domains, the RBD and the NTD (**Figure 2a**). For each of these antibodies, mutagenic libraries theoretically comprising all possible sets of mutation between the mature and inferred universal common ancestor (UCA) were constructed using combinatorial nicking mutagenesis^32,33^ and the libraries were pooled in approximately equimolar ratios and assembled into the yeast surface vector with a target of multiple barcodes per antibody variant (**Figure 2a**).

Several deep mutational scanning protocols pair a barcode to an encoded protein variant using long-read sequencing^10,34–37^. MAGMA-seq is compatible with both long-read sequencing and short-read sequencing. For short-read sequencing, the barcode is separately paired with the V_H_ and V_L_ using independent Golden Gate intramolecular ligation reactions^38^, which places the barcode adjacent to either the CDR H3 or the CDR L3 (**Figure 2b**). The reaction products are separated on an agarose gel to remove concatemers and isolate the correct intramolecular ligation product (**Extended Data Figure 3**), and amplicons are prepared for paired end short-read sequencing. PCR-based amplicon preparation of mixed populations is known to result in chimera formation between closely related nucleic acid sequences^35,39^. We evaluated several different amplicon preparation protocols by assessing chimera formation between three isogenic plasmids containing distinct mutations and unique barcodes. Using this approach, we identified a protocol resulting in low amounts of overall chimera formation (**Extended Data Figure 4**).

**Fig. 3.**
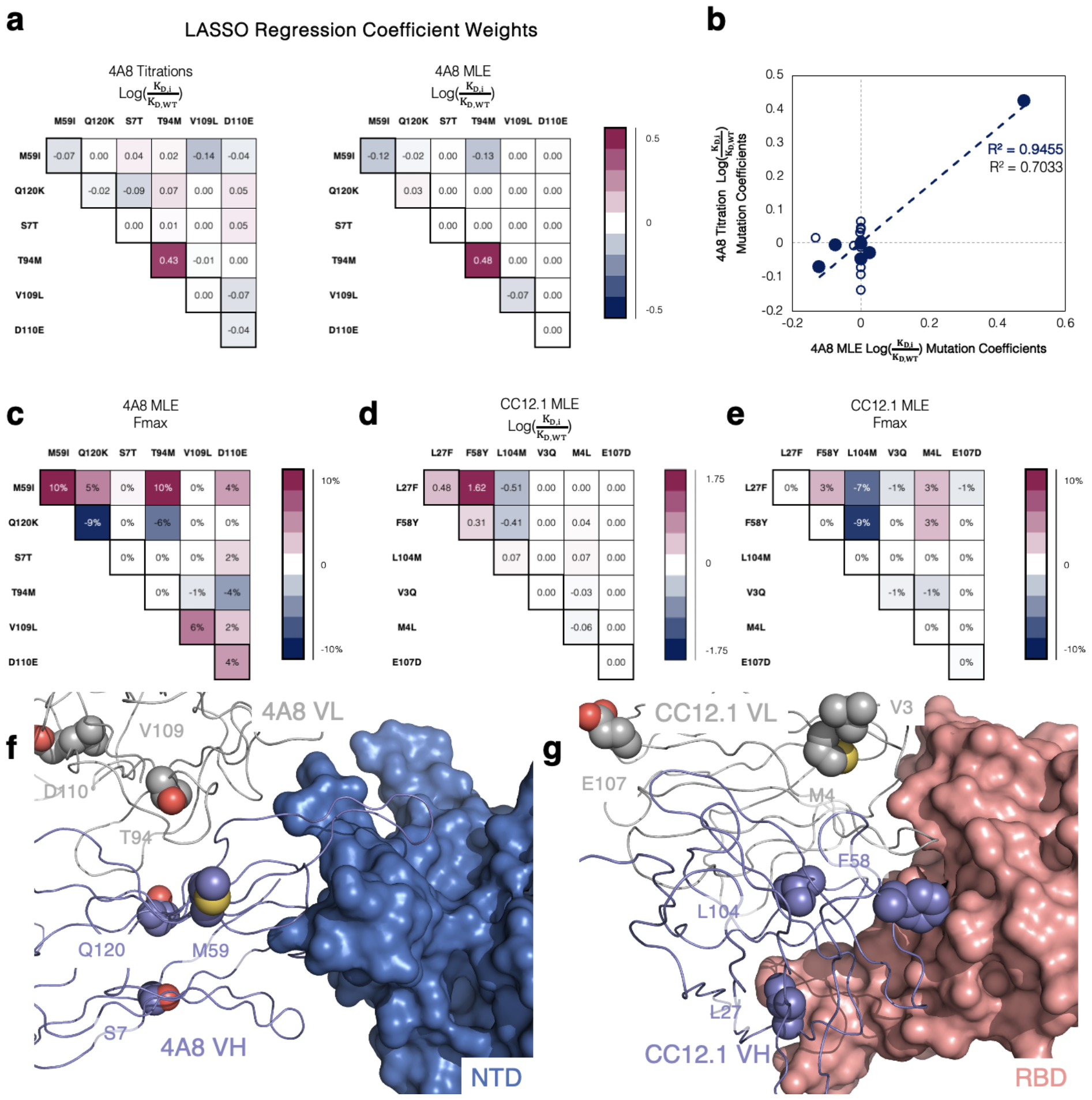
Antibody development landscapes for 4A8 and CC12.1 are sparse. (a) Comparison of 4A8 one body and two body parameter binding affinity weights inferred from (left) isogenic titrations and (right) MAGMA-seq. Binding affinities are represented as logK_D_ ratios relative to the mature antibody sequence. (b) Correlation between isogenic titrations and MAGMA-seq parameter weights. Blue closed circles are one-body weights, and open circles are two-body weights.(c) Parameter weights for 4A8 Fmax percentage differences relative to the mature antibody. (d-e) CC12.1 MLE parameter weights for (d) logK_D_ ratios and (e) Fmax as inferred from MAGMA-seq. (f-g) Structural complexes of SARS-CoV-2 Wuhan Hu-1 S antibodies (f) 4A8 bound to NTD (PDB ID: 7C2L), and (g) CC12.1 bound to RBD (PDB ID: 6XC3). Positions mutated from the inferred UCA sequence are shown as purple (V_H_) or gray (V_L_) spheres.

**Fig. 4.**
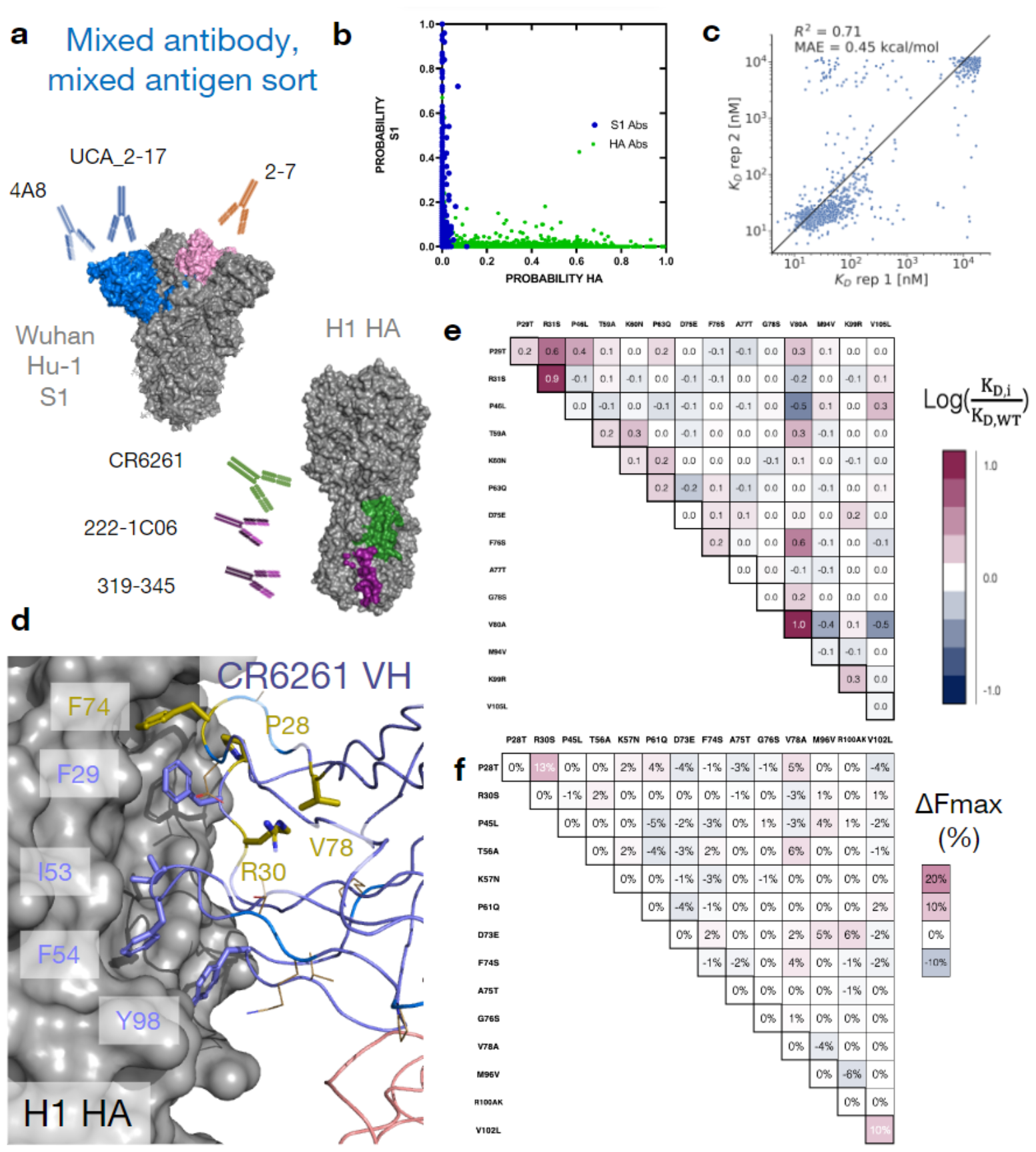
MAGMA-seq infers biophysical properties in mixed antibody, mixed antigen sorts. (a) Antigen-specific antibody sub-libraries and antigens used in the sorting experiments. Non-binders 1G01 and 1G04 were also included. (b) Probability of an antibody variant sorted into an antigen-specific bin when only 2000 nM of S1 (Y-axis) or H1 HA (X-axis) was incubated with yeast. For HA, only the top sorting bin was included in the analysis. (c) Correlation of MLE KD estimates from sort replicates for anchor epitope targeting antibodies 222-1C06 and 319-345. (d) Structure (PDB ID: 3GBN) of CR6261 bound to H1 HA. Purple sticks are HA-contacting positions that are encoded from the inferred UCA sequence. The side chains of residues mutated in the mature antibody relative to the UCA are shown as gold sticks and lines. (e-f) LASSO regression of (e) logK_D_ ratios and (f) F_max_ percentage differences one body and two body weights for CR6261. Weights are shown relative to the mature antibody.

To evaluate the fidelity of our protocol, we sequenced 20 isogenic clones using Oxford Nanopore sequencing. The pooled, mutagenic antibody library was prepared in replicates for Illumina short-read sequencing following our optimized protocol for both V_H_ and V_L_ pairings. 95% (19/20; replicate 1) and 85% (17/20; replicate 2) of barcode-antibody pairing was identical between nanopore and short read sequencing (**Figure 2c**), and no incorrect calls were made in either replicate. In total, we paired 1059 barcodes and recovered 64/64 CC12.1 variants (100% library coverage), 48/64 COV2-2489 variants (75% library coverage) after an alternative filtering step (**Extended Data Figure 5**), and 56/64 4A8 variants (87.5% library coverage) with a mean of 4.8 barcodes per variant (**Figure 2d**).

**Fig. 5.**
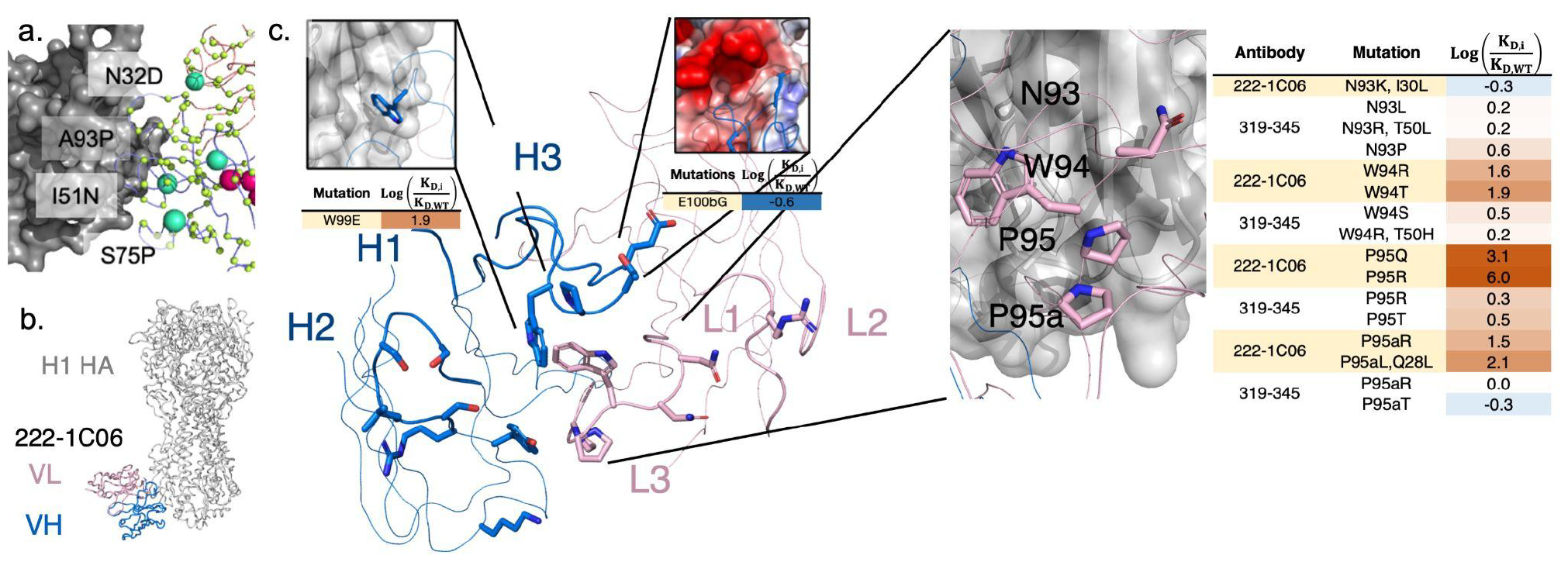
MAGMA-seq samples the function sequence-binding landscape for neutralizing antibodies. (a) Forward trajectories of the UCA of anti-S1 nAb 2-17. The sampled library is a subset of all potential single nucleotide substitutions in both VH and VL. All sampled positions are shown with CA atoms shown as lime spheres. Larger cyan spheres encode gain of function antibody variants, (b) Previously solved structure of 222-1C06 bound to H1 HA (PDB ID 7T3D). (c) 222-1C06 paratope and mutational profiles for certain residues in the CDR H3 and KL3. CDRs L1-L3 and H1-H3 are shown as larger width ribbons than the rest of the main chain. Residues with a CB within 5 A of HA are shown as colored sticks. The panel inset for the E 100bG mutation shows the electrostatic potential surface ofH1 HA.

The library was transformed into yeast, passaged, and induced by galactose. We sorted the library at 10 different S1 labeling concentrations by sorting yeast cells into two bins by fluorescence using the channel corresponding to binding S1 (**Figure 1a, Figure 2e, Extended Data Figure S6**). We sequenced and counted the number of barcodes collected from each of the bins at every sampled concentration as well as a reference population of Fab displaying cells. The count data were aggregated with fluorescence bin limits, sorted cell counts, and predetermined parameters describing the expected fluorescence distributions, and then analyzed by a custom MLE algorithm to generate monovalent binding dissociation constants (K_D_) and max mean fluorescence at saturation (F_max_) estimates for each variant. Our MLE algorithm performs minimization of the difference between observed and expected sequencing counts given an underlying system of equations describing the theoretical distributions and anticipated measurement error (for full details, see **Supporting Note 1**). Importantly, the algorithm can quantify K_D_ estimate uncertainty (**Figure 2f**). Distributions of K_D_ estimates were observed to be consistent across barcodes of the same variant, with high overlap between confidence intervals (**Figure 2g** and **Extended Data Figure S7**). Our MLE algorithm uses two fixed global parameters relating to the estimated error rate in FACS and the fluorescence probability distribution of the expressed constructs. We evaluated the sensitivity of the output on these parameters, finding that the mean absolute error in logK_D_ ratio ranged from 0.016 - 0.039 log_10_(K_D_/K_D,wt_), showing little effect overall on our parameter choices (**Extended Data Figure S8**).

To address whether parameter estimates from MLE are consistent with isogenic titrations, we used combinatorial nicking mutagenesis^32^ to prepare biological replicates for 61 separate 4A8 variants. For each variant, we performed four isogenic titrations (n=4; 2 technical replicates and 2 biological replicates of each, see **Supplementary Data 2**) and determined the change of free energy of binding upon mutation (ΔΔG) relative to the mature 4A8 Fab. While we observed a single outlier, likely because of low sequencing coverage (average counts per bin = 7, **Figure 2h**), the mean absolute error of MLE generated K_D_s relative to the isogenic titrations fell at or below the level of precision of the isogenic titrations for almost all variants tested (isogenic titration experimental limit = 0.21 logK_D_/logK_D,ref_, **Figure 2h**). Additionally, the MLE algorithm captured the statistically significant differences in F_max_ that are known to exist between 4A8 and CC12.1 Fabs from isogenic titrations (**Figure 2i**). Thus, MAGMA-seq can recover biophysically meaningful parameters that are consistent with isogenic titrations.

We performed regression analyses on the MAGMA-seq output to gain insight into the impact of individual mutations as well as to determine epistatic effects of mutations on the overall development trajectory for the 4A8, CC12.1, and COV2-2489 antibodies. As expected, due to the high K_D_ and low F_max_ observed for COV2-2489 WT (see **Figure 1c**), we noticed that few barcodes from any variants of this antibody appeared in any of the sorted bins at substantial quantities and similar analysis was not completed. For 4A8 and CC12.1, we performed one-hot encoding of the programmed mutations and then analyzed each antibody separately using different regression techniques (Ordinary Least Squares (OLS), Least Absolute Shrinkage and Selection Operator (LASSO)^40^, and Ridge Regression^41^ (**Supplementary Data 3**). While agreement was observed amongst all regression methods, we selected the LASSO due to the parameter minimization inherent to the method.

LASSO regression for the 4A8 isogenic clone logK_D_ ratio titration data and a 2^nd^ order model fit the data with MAE = 0.099 log_10_(K_D_/K_D,wt_). All 2^nd^ order coefficient weights fell below 0.07 log_10_(K_D_/K_D,wt_) (less than 17% absolute difference in binding affinities), supporting a sparse development pathway (**Figure 3a**). An identical analysis performed on the 4A8 MLE dataset reproduced the same sparse pathway results (**Figure 3a;** MAE = 0.063 log_10_(K_D_/K_D,wt_)). Surprisingly, only the light chain mutation M94T had any appreciable effect on binding. The coefficient weights for the 4A8 titrations and MLE proved consistent with a correlation coefficient of 0.94 for all first order weights (**Figure 3b**). The correlation coefficient for all first and second order weights is lower at 0.70 due to the noise present in the titration data collection (**Figure 3b**). MAGMA-seq also allowed us to perform regression analysis on F_max_, a proxy for the total amount of active Fab on the yeast surface. For 4A8, a 2^nd^ order model showed F_max_ is influenced by multiple mutations. I59M decreases Fmax by 10%, and K120Q improves F_max_ values by 9% compared to mature 4A8 (**Figure 3c**).

Analogous regression for antibody CC12.1 was performed using the MLE data for ΔΔG_binding_ and Fmax. A second order model described the data with MAE = 0.07 log_10_(K_D_/K_D,wt_) and 3923 RFU for ΔΔG_binding_ and Fmax, respectively. Consistent with 4A8, we found a sparse mutational landscape with CC12.1 and S1 where only two mutations, F27L and Y58F, are required for enhanced affinity (**Figure 3d**). M104L improves Fmax values by approx. 16% in the presence of F27L and Y58F (**Figure 3e**).

4A8 binding to S1 is mediated predominantly by the V_H_ chain with important contacts to the NTD in HCDR1 and HCDR3^29^. V_L_ M94T is the only one of six mutations from germline that improves binding affinity. A structural hypothesis for this mutation is that it repositions the HCDR3 in a more productive conformation for NTD recognition (**Figure 3f**). CC12.1 uses both V_H_ and V_L_ to contact S RBD^42^ (**Figure 3d**). Y58F directly contacts the RBD surface for improved binding, while F27L may subtly reposition the CDR H1 for improved recognition. M104L decreases binding affinity in the context of F27L and Y58F but improves functional expression, and it may participate in subtle antibody-antigen rearrangements which could cause the minor 2-body effects seen (**Figure 3d, g)**. MAGMA-seq alone as well as in combination with known structures can aid in the structural and genetic understanding of antibody development trajectories.

To determine whether MAGMA-seq can evaluate multiple antibodies sorted against multiple antigens simultaneously, we prepared a library containing mutants of eight distinct antibodies^29,43–46^ (1G01, 1G04, 319-345, 222-1C06, CR6261, 2-7, UCA_2-17, and 4A8) containing varying light chain gene usage and CDR H3 length. 1G01 and 1G04 bind at the active site on NA influenza neuraminidase N2 A/Brisbane/10/2007^43^. CR6261 is a bnAb binding to group I HAs^45^. 319-345 and 222-1C06 are nAbs which recognize the anchor epitope on H1 HA^44^. 2-7, UCA_2-17, and 4A8 recognize SARS-CoV-2 spike Wuhan Hu-1^29,46^ (**Figure 4a, Figure 1C**). We sorted replicates of this library of 4,105 matched barcoded antibodies against 11 varying combined concentrations of HA and S1. The 11 sorts were structured such that, at all labeling concentrations, the average population had an appreciable binding signal (**Extended Data Figure S9**). One labeling concentration contained only HA or S1, respectively. Additionally, the library contained internal controls for evaluating the sorting error and for assessing the fidelity of affinity reconstruction. The complete dataset for all antibody variants is listed in **Supplementary Data 4**. As expected, none of the NA-specific 1G01 or 1G04 antibody variants had inferred dissociation constants below 1 μM for either the HA or S1 antigen. HA-specific and S1-specific antibodies mapped neatly to one of the two antigens using the antigen-only sort (**Fig 4b**). The 4A8 variants, included as internal controls, were consistent with the parameter weights from the previous sort (LogK_D_ ratio of T94M relative to the S7T variant: 1.33). Additionally, the estimated K_D_ values from MLE are reasonably consistent between replicate sorts. After removing variants containing stop codons and non-converged values, we observe an R^2^ of 0.71 and MAE of 0.32 log_10_(K_D_/K_D,wt_) for anchor antibodies 222-1C06 and 319-345 (**Fig 4c)**. Relative to replicate 1, replicate 2 underpredicts some of the intermediate affinity antibodies. We attribute this discrepancy to the absence of the 100nM labeling bin for the second replicate.

Two antibodies in the library contained mutations allowing for the reconstruction of potential development trajectories from their inferred UCA sequence. 2-7 is a Wuhan Hu-1 S1-specific nAb^46^. 2-7 contains five mutations from its inferred germline, all in the V_L_ (A5G, A31G, D52E, K55N, T95S). The inferred development (**Extended Data Figure S10**) was superficially like the sparse development pathways observed for 4A8 and CC12.1, with four 1^st^ order couplings predicting the dissociation constants of the potential pathway variants, as supported from LASSO regression.

CR6261 is an influenza bnAb targeting the HA stem epitope originally described by Throsby et al^45^. It is an unusual antibody in two ways. First, its development trajectory is dissimilar to other VH1-69 anti-HA antibodies previously characterized^47^. Second, it confers molecular recognition only through its V_H_, mainly by positioning apolar residues at framework 3 (FR3) (F74, V78), in CDR H1 (F29), and in CDR H2 (F54) in a hydrophobic groove^48,49^ (**Fig 4d**). Both CDR residues are encoded in germline VH1-69 sequence in allelic human populations, but the inferred UCA sequence does not appreciably recognize the H1 HA stem epitope^50^. The potential first steps of its trajectory from its inferred UCA sequence have been developed by Lingwood et al.^50^, supporting a first committed step of some combination of H1 mutations T28P and S30R necessary for orientation of F29, and at least some subset of the framework 3 (FR3) mutations (E73D/T75A/S76G/A78V) necessary for F74 insertion (**Fig 4d**). We sampled 2.9% (470/16,384 possible variants) of the potential sequences between the UCA and mature CR6261. MAGMA-seq recovered a K_D_ of 12 nM for mature CR6261, consistent with isogenic titration of 9.4 nM and with previous literature reports^45^. LASSO regression supported a 2^nd^ order epistatic model (**Supplementary Data 3)**, with a total of 5 1^st^ order and 15 2^nd^ order weights above an absolute 0.18 log_10_(K_D_/K_D,wt_) energetic threshold. Consistent with the studies from Lingwood and Pappas, the strongest 1^st^ order weights contributing to binding affinity are T28P, S30R, and FR3 mutation A80V (**Fig 4e**), and the two strongest 2^nd^ order weights are the epistatic couplings between T28P/S30R (0.65 log_10_(K_D_/K_D,wt_)) and S74F/A80V (0.57 log_10_(K_D_/K_D,wt_)). The known epistasis in the T28P/S30R mutations can be rationalized as altering the orientation of the CDR H1 loop such that F29, usually buried, largely becomes solvent exposed in the unbound structure. Consistent with this hypothesis, the surface expression of Fabs containing T28P/S30R mutations decreased by approximately 10% (**Fig 4f**), as expected for mutations which increase the apolar solvent accessible surface area. The other epistatic relationship observed of S74F/A78V can relate to the positioning of hydrophobic residues, where the 78V is needed to constrain the correct F74 rotamer for precise shape complementarity in the stem groove. In sum, the sparse sampling of bnAb mutants allow for the reconstruction of the development pathways that are in concordance with the existing body of structural, genetic, and immunological evidence for this antibody. Thus, MAGMA-seq can reconstruct the likely development pathways for multiple human antibodies against different antigens in the same experiment.

The libraries described thus far are all retrospective analyses of antibody development trajectories, where libraries encoded chimeras of the mature and UCA sequences. To further investigate the utility of this method, our second demonstration of MAGMA-seq included a prospective antibody development library and a few CDR targeted site-saturation mutagenesis libraries. We generated each of these antibody libraries in parallel reactions and subsequently pooled and barcoded the variants. We bottlenecked the library, which selected individual variants randomly, and assessed it with MAGMA-seq.

To test whether MAGMA-seq could map prospective antibody development trajectories, the mixed library contained a subset of a larger library of the UCA sequence of the anti-S NTD 2-17 (UCA_2-17)^46^. This larger library theoretically contained all single nucleotide substitutions at the CDRs and framework positions. Its UCA was predicted to bind at a K_d_ of 2050 nM (range 400-3200 nM; 0.23 log_10_(K_D_/K_D,wt_) s.d.; 58 barcoded UCA sequences) and a mean F_max_ = 890, consistent with measurements of the isogenic control (**Fig 1C**). We were able to recover 318 uniquely barcoded variants. Many of these mutants, like V_H_:Y91DHN or V_H_:C92GFY near the CDR H3, are expected to structurally destabilize the protein, resulting in non-specific binders. Still, several mutants had lower inferred dissociation constants or higher F_max_ values than the UCA, including V_H_:I51N in CDRH2 (K_d_ 970 nM) and V_L_:N32D (F_max_ 4,000) observed in the mature 2-17 sequence, V_H_:S75P (K_d_ 400 nM), and V_H_:A97P in CDRH3 (K_d_ 490 nM) (**Fig 5a**). Thus, MAGMA-seq can evaluate potential forward trajectories for antibodies that are consistent with genetic and structural data.

We also used MAGMA-seq to infer the preliminary rules of recognition for an emerging class of influenza neutralizing antibodies. Antibodies 319-345 and 222-1C06 target a distinct anchor stem epitope of H1 HA^44^. Anchor bnAbs appear to be germline restricted to light chains VK3-11 or VK3-15, with heavy chains from germlines VH3-23, VH3-30/VH3-30-3, and VH3-48. All mature anchor bnAbs encode a CDR H3 of diverse amino acid sequences, with a glycine either at the beginning or end of the CDR H3 and two to four hydrophobic residues at the middle of the sequence. The cryo-EM structure of 222-1C06 bound to H1 HA shows the structural basis of recognition. The interaction at the anchor epitope is dominated by multiple hydrophobic interactions across the heavy and light chains. The germline-encoded and invariant CDR KL3 ‘NWPP’ motif from positions 93-95A are at the center of the binding interface. CDRH2 (Leu55) and CDRH3 (Trp99, Pro100, Thr100a) all contribute hydrophobic contacts at the binding interface (**Fig 5b**,**c**).

We recovered 183 and 390 single non-synonymous mutants of 222-1C06 and 319-345, respectively (1429 uniquely barcoded variants). The observed K_D_ for mature antibodies were low nM (319-345: 16 nM; 222-1C06: 27 nM) and highly reproducible between independent barcodes (319-345: 0.092 log_10_(K_D_/K_D,wt_) s.d., n=171; 222-1C06: 0.07 log_10_(K_D_/K_D,wt_) s.d., n=92). CDR loops L1, L2, and H1 make peripheral contacts at the interface. Consistent with this, only 3.8% of single mutants (2/118 and 11/161 for 222-1C06 and 319-345, respectively) at CDR L1, L2, and H1 positions disrupted binding affinity by greater than 0.7 log_10_(K_D_/K_D,wt_) (**Supporting Data 4)**. This contrasts with CDR H2, where 40% (20 of 51) of single and double mutants disrupted binding greater than 0.7 log_10_(K_D_/K_D,wt_), supporting the importance of H2 in recognition of the anchor epitope (**Fig 5c**). While the library under sampled CDRH3, mutations at Trp99 for 222-1C06 (W99E log(K_D,i_/K_D,WT_) 1.9) and Gly100d for 319-345 (G100dL/I >2.1 log_10_(K_D_/K_D,wt_)) were deleterious, consistent with the precise positioning of the loop needed for binding. In the KL3 ‘NWPP motif’, observed mutations at N93 seem to have little effect on binding affinity, while mutations at W94, P95, and P95a seem to drastically disrupt binding in 222-1C06 (**Fig 5c**). Intriguingly, mutations at these same positions in 319-345 are only mildly deleterious (**Fig 5c**), suggesting that the antibody paratopes are positioned slightly differently against HA.

To identify candidate mutants with lower binding affinities than the mature antibodies, we identified all variants with log(Kdi/Kd,wt) values falling at least two standard deviations below zero. No 319-345 mutants met this cutoff, while four 222-1C06 variants did (VH:E100bG, VH:S54G, VH:D101G, and VH:D101S; **Fig 5c**). E100b is adjacent to an acidic patch on HA in the structural complex (**Fig 5c**), and so mutation to glycine likely improves binding by eliminating this unfavorable electrostatic contact. The mechanistic basis of the D101 mutations remains unclear, as mutation likely disrupts a salt bridge with CDRH3 R94. Likewise, the effect of S54G is obscure, although we note that this mutation occurs in several 319-345 clonotypes^44^ isolated from patients.

## Discussion

In this paper we present MAGMA-seq, an integrated technology for quantitative wide mutational scanning of human antibody Fab libraries. We demonstrate MAGMA-seq on two pooled libraries comprising mutants of ten different human antibodies spanning light chain gene usage, CDR H3 length, and antigenic targets. Analysis of MAGMA-seq outputs allows for the simultaneous mapping of retrospective and prospective potential antibody development pathways, paratope affinity maturation, and the sequence dependence on binding for broadly neutralizing antibodies. MAGMA-seq can be deployed immediately not only in these areas but for affinity maturation campaigns, specificity mapping campaigns, and for fine paratope mapping. A compelling advantage of MAGMA-seq is its ability to measure binding for mutants of many given parental antibodies in a single experiment. Since modern biotech campaigns typically use dozens of candidates in initial testing, MAGMA-seq enables the streamlining of such measurements.

We used MAGMA-seq to reconstruct potential development pathways for anti-influenza (CR6261) and anti-SARS-CoV-2 (4A8, CC12.1, 2-7) nAbs. We found that these development pathways can be reconstructed by considering binding contributions from only a handful of the mutations. This is supported by a body of evidence from other protein families^51,52^ showing the sparseness of functional protein landscapes^53^. We also found that these sequence-binding fitness landscapes were most consistent with one-body or at most two-body interactions, consistent with recent protein engineering literature^54–56^. The resulting implication is that sampling of a small percentage of potential variants is sufficient for reconstruction of fitness landscapes. Indeed, for the CR6261 experiments we sampled 470 out of 16,384 possible variants and were still able to reconstruct a development trajectory supported by existing evidence. Likely many such antibody trajectories can be inferred from relatively few experiments.

We also evaluated the sequence dependence of two newly described nAbs targeting the anchor epitope on influenza HA. Our broad findings established the importance of several key mutations at the antibody side of the interface, identified electrostatic complementarity as a mechanism for improving nAb recognition to the anchor epitope, and highlighted the importance of shape complementarity for the diverse CDR H3 sequences found to fit in the interface. We anticipate that MAGMA-seq will be used to enumerate the sequence determinants for entire sets of antibodies targeting key neutralizing or other important antigenic epitopes.

There are some limitations with the current demonstration of this technology. First, we assess binding using yeast surface display, limiting the practical dynamic range of binding affinities to 0.5 nM - 2 μM. At high affinities, the labeling time to reach equilibrium reaches >10 hours, and at low affinities the antigen can dissociate off the yeast surface during sorting^57^. Many therapeutic antibodies with low picomolar monovalent binding affinities would be impracticable to assess accurately. Second, we have measured order of magnitude differences in F_max_ values between different mature Fabs (see **Fig 1c**). Evidence suggests some correlation between functional expression on the yeast surface and stability of variants deriving from the same parental sequence^6,58^, but a complete understanding of what drives differential Fab expression *between* parental Fabs is not yet known. The low F_max_ values of some antibodies can hinder MLE performance solely due to the variant having low probability of being sorted into a bin, which was exemplified by the low counts of antibody COV2-2489 variants collected in our first demonstration. Third, the MLE algorithm uses one global parameter that cannot be measured during the experiment. Despite this limiting assumption, the inferred monovalent dissociation constants match published results where known. Fourth, no explicit removal of non-specific binders, like that seen for the anti-S NTD 2-17 sorts, were performed here. A parallel sort with polyspecificity reagent could improve discrimination of bona fide binders. Fifth, we note that, due to the implementation of FACS with yeast display, accuracy of MAGMA-seq estimated binding affinities may not precisely match gold-standard in vitro measurements like SPR, where antibody/antigen interactions are more directly quantified. Additional encumbrances to the method presented include the formation of antibody sequence chimeras during intramolecular ligation that reduce the number of identified barcodes and the use of TruSeq small RNA single 6-nt index adapters that allow for more index hopping during Illumina sequencing. Technical improvements would remain compatible with the rest of the MAGMA-seq workflow. Long-read sequencing is becoming increasingly inexpensive and more accurate, and as it improves it removes the necessity of PCR amplification.

We demonstrated this technology on libraries of fewer than 10,000 variants, although the functional limit on the library size is much larger. The potential complexity bottlenecks for library size are through generation of individual mutagenic libraries, Gibson assembly into barcoded yeast display plasmids, transformation into yeast, sorting in yeast, and sequencing. An additional complexity bottleneck arises through the linking VH and VL genotypes via barcodes. The major bottleneck at the current stage of development is through cell sorting. For sorting speeds of commercially available cell sorters, the protocol leads to approx. 1,000 cells collected per sorting bin (10,000 events per second x 40% Fab displaying cells per event x 25% collection of the Fab displayed cells). Since we sample at least 150-fold above the theoretical size of the library, this means that a library size of 10,000 would take 25 minutes per labeling concentration. Sorting the full suite of 10-12 labeling concentrations would then take a full working day, including start-up and shutdown. Significantly larger libraries would require multiple days of sorting or multiple cell sorters running in parallel.

## Outlook

Massively parallel measurements of protein binding affinities can be used to train deep learning models to capture antibody molecular recognition. We have demonstrated that this MAGMA-seq technology can perform wide mutational scanning for multiple antibodies against different antigens over a wide dynamic range of binding affinities. These measurements are made in a natural human Fab background and have multiple internal controls needed for quality control and validation. The next steps are an integrated computational and experimental appraisal of the quality and quantity of data needed for such purposes.

## Methods

### Materials

All media components were purchased from ThermoFisher or VWR. All enzymes were purchased from New England Biolabs unless otherwise specified. The recombinant SARS-CoV-2 Spike S1-hFc-His tagged protein used for titrations and sorting was purchased from ThermoFisher (RP-876-79). The recombinant neuraminidase (NA) for titrations was obtained through BEI Resources, NIAID, NIH: N2 Neuraminidase (NA) Protein with N-Terminal Histidine Tag from Influenza Virus, A/Brisbane/10/2007 (H3N2), Recombinant from Baculovirus, NR-43784. The ectodomain of A/Brisbane/02/2018 H1 HA with a foldon trimerization domain was expressed in HEK293T cells (ATCC) and purified using Ni-NTA affinity chromatography. Recombinant neuraminidase and recombinant hemagglutinin were biotinylated in a 20:1 molar ratio of biotin to antigen with EZ-Link NHS-Biotin (ThermoFisher, 20217) following the manufacturer’s instructions.

### Plasmids

All plasmids were constructed using either NEBuilder HiFi DNA Assembly Master Mix (New England Biolabs) for Gibson assembly^28^, by Golden Gate assembly^27,59^, using a Q5 Site-Directed Mutagenesis Kit (New England Biolabs), or by nicking mutagenesis^32,60^. Synthetic DNA was ordered either as gBlocks or eBlocks (IDT). A complete list of plasmids, libraries, gene blocks, and primers are located in **Supplementary Data 1**.

### Construction of Fab libraries

Fabs were diversified either by complete combinatorial mutagenesis^32^, site-saturation mutagenesis^61^, or oligo pool nicking mutagenesis^62^. Complete combinatorial libraries of Fabs were prepared from mature human antibodies and their inferred universal common ancestor (UCA). UCA sequences were inferred using IgBLAST^63^. In total, 10 mutagenic libraries were prepared (all library details are in **Supplementary Data 1**). Fab libraries were combined with barcoded yeast display plasmid(s) by Gibson assembly^28^ and bottlenecked. Five μg of plasmid DNA was transformed into chemically competent *Saccharomyces cerevisiae* (EBY100, ATCC MYA-4941) and stored as yeast glycerol stocks in -80 °C according to Medina-Cucurella & Whitehead^64^.

### Barcode-variant pairing

Barcodes were paired with V_H_ and V_L_ variants through Oxford nanopore sequencing or by short-read sequencing of amplicons prepared by intramolecular ligation of barcode in proximity to the CDR3 of either the V_H_ or V_L_ using Golden Gate^27^. Oxford nanopore sequencing (Plasmidsaurus) was performed on individual plasmids. Short-read amplicons were sequenced on an Illumina MiSeq with 2x250 paired end reads (Rush University Sequencing Core). For intramolecular ligation, two replicates were performed independently.

### Yeast Cell Surface Titrations

To determine the binding affinity of individual variants, isogenic titrations were performed according to Chao et al.^65^. 4A8 variants were made by the method of combinatorial nicking mutagenesis^32^. Each variant was tested in duplicate on two separate days (n=4 total replicates) and compared with a titration of mature 4A8 Fab to determine ΔΔG_binding_, the free energy of binding upon mutation. The isogenic titrations reported in **Figure 1** were reported in at least duplicate (n≥2).

### Sorting of Fab libraries

For sorting the mixed 4A8/COV2-2489/CC12.1 library, 1e7 (ten million) yeast library cells from glycerol stocks were shaken at 230 rpm and grown in 250mL flasks at 30 °C overnight in 50 mL SDCAA + PenStrep and kanamycin. The next day, the 1e7 yeast cells were induced in SGDCAA + PenStrep and kanamycin at 20 °C for 48 hours in a total reaction volume of 50 mL. On the morning of sorting the cells were concentrated to an OD_600_ = 5 in ice-cold PBSF. Ten million library cells were then labeled with different amounts of S1-hFc-His at the following concentrations in nM: 0, 1, 2.5, 5, 10, 50, 100, 250, 500, 1000, 2000 for 30 minutes at room temperature. After the binding reactions were finished cells were spun down, washed with 1mL of ice-cold PBSF, and then labeled with 6.25 μL anti-V5-AlexaFluor488 and 25 μL Goat anti-hFc-PE (ThermoFisher, 12-4998-82) for 30 minutes covered on ice. After fluorophore labeling, the cells were pelleted and washed with 1 mL of ice-cold PBSF, and pellets were left covered on ice until loading onto Sony SH800 cell sorter, at which time each pellet was resuspended in 5 mL of ice-cold PBSF. Cells were first gated for yeast cells and single cells (drawn according to Banach et al.^66^ to avoid collection of clumped yeast of irregular large yeast aggregates), and then a gate for positive Fab expression was drawn and 200,000 cells were collected as the library reference population (**Extended Data Figure 6**). Sorting bins for the Top 25% and Next 25% of binding based on PE signal were gated from the display positive population and 200,000 cells were collected in each bin (**Extended Data Figure 6**). Sorted cells were recovered in 1 mL of SDCAA plus antibiotics overnight at 30 °C, at which time another 1 mL of SDCAA was added. Cells were grown until they reached an OD_600_ greater than 2. Cell stocks were made for each sorted population at 1 mL of OD_600_ = 1 in yeast storage buffer (20% w/v glycerol, 20 mM HEPES-NaOH, 200 mM NaCl, pH = 7.5).

For sorting the S1/HA library, 1e7 (ten million) yeast library cells from glycerol stocks were shaken at 230 rpm and grown in 250mL flasks at 30 °C overnight in 50 mL SDCAA + PenStrep and kanamycin. The next day, the 1e7 yeast cells were induced in SGDCAA + PenStrep and kanamycin at 20 °C for 48 hours in a total reaction volume of 50 mL. On the morning of sorting the cells were concentrated to an OD_600_ = 5 in ice-cold PBSF. Ten million library cells were then labeled with different amounts of S1-hFc-His and biotinylated HA for 30 minutes at room temperature. The 11 labeling concentrations spanned from 2.5 nM – 2000 nM and included mixes of both S1-hFc-His and biotinylated HA. After the binding reactions were finished cells were spun down, washed with 1mL of ice-cold PBSF, and then labeled with 6.25 μL anti-V5-AlexaFluor488, 25 μL Goat anti-hFc-PE (ThermoFisher, 12-4998-82), and 25 μL SAPE (ThermoFisher, S866) for 30 minutes covered on ice. After fluorophore labeling, the cells were pelleted and washed with 1 mL of ice-cold PBSF, and pellets were left covered on ice until loading onto Sony SH800 cell sorter. Each pellet was resuspended in 5 mL of ice-cold PBSF and loaded on to the cell sorter. Cells were first gated for yeast cells and single cells, and then a gate for positive Fab expression was drawn and 1,000,000 cells were collected per bin for the first replicate and 750,000 cells per bin were collected for the second replicate. Sorted cells were recovered in 5 mL of SDCAA plus antibiotics for at least 30 hours at 30 °C and cell stocks were made for each sorted population in yeast storage buffer. Yeast biological replicates were performed. The plasmid encoded master library was prepared once and separately transformed into yeast; these libraries were sorted on separate days.

### Amplicon Preparation and Deep Sequencing

Plasmid DNA from each collected population was prepared according to Medina-Cucurella & Whitehead^64^ using Zymoprep Yeast Plasmid Miniprep kits in either individual Eppendorf tubes (D2004) or 96-well plate format (D2007) and plasmid DNA was eluted in 30 μL nuclease free water. 15 μL of eluted plasmid DNA was further purified with exonuclease I and lambda exonuclease. The barcode region of the purified DNA was amplified using 25 PCR cycles with Illumina TruSeq small RNA primers following Kowalsky et al ‘Method B’^67^. Amplicons were sequenced on either an Illumina MiSeq (4A8/CC12.1/COV2-2489 sort) or NovaSeq6000 (S1/HA sorts) by Rush University with single end reads.

### Parameter Estimation

A complete description of the mathematics behind parameter estimation is detailed in **Supporting Note 1** and a description of the computational pipeline is described in the **Extended Materials and Methods**. Custom Python software was used to estimate variant-specific monovalent binding dissociation constants (K_d,i_) and mean maximum fluorescence at saturation (F_max,i_) fit by equation (1). These values were inferred using maximum likelihood estimation of the following expression for the log likelihood *LL*_*i*_*(K*_*d,i*_, *F*_*max,i*_):

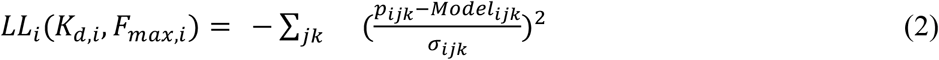

Here, p_ijk_ is the probability of capturing variant i in bin j at labeling concentration k and is determined from observables from the deep sequencing experiment according to the following equation:

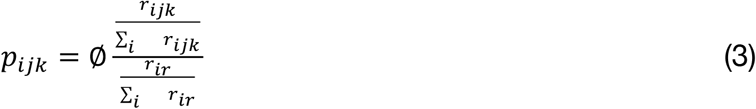

∅ is the total fraction of cells collected in the sorting bin relative to the reference sample, r_ijk_ is the number of observed read counts for variant i in bin j at labeling concentration k, r_ir_ is the number of observed read counts for variant i in the reference population, and the summations represent the sum of observed read counts over all barcodes.

*Model*_*ijk*_ is the model probability of the variant i sorting in bin j at labeling concentration k and is defined as:

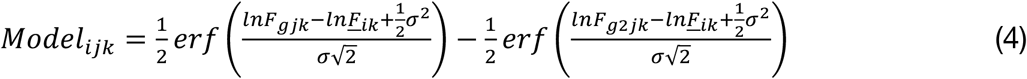

Here, *F*_*gjk*_ *and F*_*g*2*jk*_ are the gating boundaries in the selected bin j, and σ is the standard deviation of the log normal distribution and set to 1.02 for all variants. Different parameter values in equation (1) change the variant-specific mean fluorescence *<u>F</u>*_*i*k_ at each labeling concentration used in the experiment.

The parameter σ_*ijk*_ representing the uncertainty in the probability of sorting is defined as:

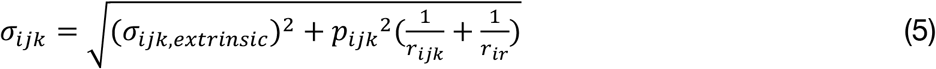

For sorts reported in Figure 2, σ_*ijk,extrinsic*_ was *set to* 0.02. For the sorts reported in Figure 4, this value was measured using the average probabilities of the non binding mutants of antibodies 1G01 and 1G04.

### Supervised Learning

Programmed mutations for reverse trajectory libraries were one-hot encoded using the custom python notebook One-hot-encode.ipynb. Ordinary Least Squares (OLS), Least Absolute Shrinkage and Selection Operator (LASSO), and Ridge Regression analyses were performed on the one-hot encoded variants for log(K_D,i_/K_D,WT_) (4A8 titrations and MLE) and Fmax (MLE) regularization using custom python jupyter-notebooks OLS.ipynb, LASSO.ipynb, and Ridge.ipynb. Coefficient weights and error values for each regression technique and model order are detailed in **Supplementary Data 3**.

Sequences of anchor mAbs used in this study are from Guthmiller et al.^44^ Clonal analyses were performed using VGenes (https://wilsonimmunologylab.github.io/VGenes/) using sequences from Guthmiller et al.

## Reporting Summary

Further information on research design is available in the Nature Research Reporting Summary linked to this article.

## Data availability

Processed deep sequencing data is available on sequencing read archive (SRA Deposition #s to be added upon publication). The plasmids for constructing compatible workflow Fabs pBDP, pMMP_kappa, pMMP_lambda, pYSD_kappa_mRFP, and pYSD_lambda_mRFP, as well as positive control plasmids p4A8_S7T_BC and p4A8_M59I_T94M_BC, are freely available from AddGene; numbers to be added upon publication.

## Code availability

All custom scripts and code are freely available on GitHub (https://github.com/WhiteheadGroup/MAGMA-seq).

## Acknowledgements

This work was supported by the National Institute Of Allergy And Infectious Diseases of the National Institutes of Health (Award Numbers 5R01AI141452-05 to T.A.W.; R00AI159136 to J.J.G.), the US Department of Education (Award Number P200A180034, participant support B.M.P.), the National Science Foundation Graduate Research Fellowship Program (Z.T.B. DGE Award Number 2040434, fellow ID: 2021324468), and the NSF REU (Award #2244288 for K.M.C.). This work utilized the Alpine high performance computing resource at the University of Colorado Boulder. Alpine is jointly funded by the University of Colorado Boulder, the University of Colorado Anschutz, Colorado State University, and the National Science Foundation (award 2201538). The authors also acknowledge Dan Schwartz for useful discussions around MLE, Pete Tessier around barcode tagging, John Jumper for helping coin the term ‘wide mutational scanning’, and Kevin Kunstman, Cecilia Chau, and Ashley Wu at Rush University for helpful discussions around NGS.

## Author contributions

Conceptualization: B.M.P., M.B.K., T.A.W. Designed plasmid sets: B.M.P., M.B.K., O.M.I., Z.T.B., E.R.R., T.A.W., Designed bench research: B.M.P., M.B.K., I.S., T.A.W. Performed bench research: B.M.P., M.B.K., O.M.I., I.S., C.M.H., A.M.W., S.A.U. Developed computational algorithms: B.M.P., M.B.K., K.M.C., P.J.S., T.A.W. Developed novel code: B.M.P., M.B.K., K.M.C., P.J.S. Data analysis: B.M.P., M.B.K., O.M.I., J.J.G., T.A.W. Contributed novel reagents: E.A., J.J.G. Writing: B.M.P., M.B.K., O.M.I., T.A.W. Supervision: T.A.W. Funding Acquisition: T.A.W., J.J.G., Z.T.B.

## References

1. Jumper, J. et al. Highly accurate protein structure prediction with AlphaFold. Nature 596, 583–589 (2021).

2. Brennan Abanades 1, Wing Ki Wong 2, Fergus Boyles1, Guy Georges2, A. B. 2 & C. M. D. ImmuneBuilder: Deep-Learning models for predicting the structures of immune proteins Brennan. Commun. Biol. 1–8 (2023). doi:10.1103/physics.15.181

3. Ruffolo, J. A., Sulam, J. & Gray, J. J. Antibody structure prediction using interpretable deep learning. Patterns 3, 100406 (2022).

4. Pittala, S. & Bailey-Kellogg, C. Learning context-aware structural representations to predict antigen and antibody binding interfaces. Bioinformatics 36, 3996–4003 (2020).

5. Hie, B. L. et al. Efficient evolution of human antibodies from general protein language models and sequence information alone. bioRxiv 2022.04.10.487811 (2022). doi:10.1038/s41587-023-01763-2

6. Makowski, E. K. et al. Co-optimization of therapeutic antibody affinity and specificity using machine learning models that generalize to novel mutational space. Nat. Commun. 13, (2022).

7. Prihoda, D. et al. BioPhi: A platform for antibody design, humanization, and humanness evaluation based on natural antibody repertoires and deep learning. MAbs 14, (2022).

8. Hummer, A. M., Abanades, B. & Deane, C. M. Advances in computational structure-based antibody design. Curr. Opin. Struct. Biol. 74, 102379 (2022).

9. 9. Hummer, A. M., Schneider, C., Chinery, L. & Charlotte, M. Investigating the Volume and Diversity of Data Needed for Generalizable Antibody-Antigen Δ Δ G Prediction. 1–16 (2023).

10. Wrenbeck, E. E., Faber, M. S. & Whitehead, T. A. Deep sequencing methods for protein engineering and design. Curr. Opin. Struct. Biol. 45, 36–44 (2017).

11. Phillips, A. M. et al. Binding affinity landscapes constrain the evolution of broadly neutralizing antiinfluenza antibodies. Elife 10, 1–40 (2021).

12. Kowalsky, C. A. et al. Rapid fine conformational epitope mapping using comprehensive mutagenesis and deep sequencing. J. Biol. Chem. 290, 26457–26470 (2015).

13. Kowalsky, C. A. & Whitehead, T. A. Determination of binding affinity upon mutation for type I dockerin–cohesin complexes from Clostridium thermocellum and Clostridium cellulolyticum using deep sequencing. Proteins Struct. Funct. Bioinforma. 84, 1914–1928 (2016).

14. Adams, R. M., Mora, T., Walczak, A. M. & Kinney, J. B. Measuring the sequence-affinity landscape of antibodies with massively parallel titration curves. Elife 5, 1–27 (2016).

15. Phillips, A. M. et al. Hierarchical sequence-affinity landscapes shape the evolution of breadth in an anti-influenza receptor binding site antibody. Elife 12, 1–31 (2023).

16. Sivelle, C. et al. Fab is the most efficient format to express functional antibodies by yeast surface display. MAbs 10, 720–729 (2018).

17. Mason, D. M. et al. High-throughput antibody engineering in mammalian cells by CRISPR/Cas9-mediated homology-directed mutagenesis. Nucleic Acids Res. 46, 7436–7449 (2018).

18. Goike, J. et al. Synthetic repertoires derived from convalescent COVID-19 patients enable discovery of SARS-CoV-2 neutralizing antibodies and a novel quaternary binding modality. bioRxiv 2021.04.07.438849 (2021). doi:10.1101/2021.04.07.438849

19. Shiakolas, A. R. et al. Efficient discovery of SARS-CoV-2-neutralizing antibodies via B cell receptor sequencing and ligand blocking. Nat. Biotechnol. 40, 1270–1275 (2022).

20. Rosowski, S. et al. A novel one-step approach for the construction of yeast surface display Fab antibody libraries. Microb. Cell Fact. 17, 1–11 (2018).

21. Weaver-Feldhaus, J. M. et al. Yeast mating for combinatorial Fab library generation and surface display. FEBS Lett. 564, 24–34 (2004).

22. Schröter, C. et al. A generic approach to engineer antibody pH-switches using combinatorial histidine scanning libraries and yeast display. MAbs 7, 138–151 (2015).

23. Lou, J. et al. Affinity maturation of human botulinum neurotoxin antibodies by light chain shuffling via yeast mating. Protein Eng. Des. Sel. 23, 311–319 (2010).

24. Mei, M. et al. Prompting Fab Yeast Surface Display Efficiency by ER Retention and Molecular Chaperon Co-expression. Front. Bioeng. Biotechnol. 7, 1–11 (2019).

25. Roth, L. et al. Facile generation of antibody heavy and light chain diversities for yeast surface display by Golden Gate Cloning. Biol. Chem. 400, (2018).

26. Chockalingam, K., Peng, Z., Vuong, C. N., Berghman, L. R. & Chen, Z. Golden Gate assembly with a bi-directional promoter (GBid): A simple, scalable method for phage display Fab library creation. Sci. Rep. 10, 1–14 (2020).

27. Engler, C., Kandzia, R. & Marillonnet, S. A one pot, one step, precision cloning method with high throughput capability. PLoS One 3, (2008).

28. Gibson, D. G. et al. Enzymatic assembly of DNA molecules up to several hundred kilobases. Nat. Methods 6, 343–345 (2009).

29. Chi, X. et al. A neutralizing human antibody binds to the N-terminal domain of the Spike protein of SARS-CoV-2. Science (80-.). 369, 650–655 (2020).

30. Thomas F. Rogers1, 2*, Fangzhu Zhao1, 3, 4*, Deli Huang1*, Nathan Beutler1*, Alison Burns1, 3, 4 et al. Isolation of potent SARS-CoV-2 neutralizing antibodies and protection from disease in a small animal model. Science (80-.). 369, 956–963 (2020).

31. Suryadevara, N. et al. Neutralizing and protective human monoclonal antibodies recognizing the Nterminal domain of the SARS-CoV-2 spike protein. Cell 184, 2316–2331.e15 (2021).

32. Kirby, M. B., Medina-Cucurella, A. V., Baumer, Z. T. & Whitehead, T. A. Optimization of multi-site nicking mutagenesis for generation of large, user-defined combinatorial libraries. Protein Eng. Des. Sel. 34, 1–10 (2021).

33. Kirby, M. B. & Whitehead, T. A. Facile Assembly of Combinatorial Mutagenesis Libraries Using Nicking Mutagenesis. Methods Mol. Biol. 2461, 85–109 (2022).

34. Lee, J. M. et al. Deep mutational scanning of hemagglutinin helps predict evolutionary fates of human H3N2 influenza variants. Proc. Natl. Acad. Sci. U. S. A. 115, E8276–E8285 (2018).

35. Levin, I. et al. Accurate profiling of full-length Fv in highly homologous antibody libraries using UMI tagged short reads. 1–15 (2023).

36. Starr, T. N. et al. Deep Mutational Scanning of SARS-CoV-2 Receptor Binding Domain Reveals Constraints on Folding and ACE2 Binding. Cell 182, 1295–1310.e20 (2020).

37. Kim, D. S. et al. Three-dimensional structure-guided evolution of a ribosome with tethered subunits. Nat. Chem. Biol. 18, 990–998 (2022).

38. Strawn, I. K., Steiner, P. J., Newton, M. S. & Whitehead, T. A. A method for generating user-defined circular single-stranded DNA from plasmid DNA using Golden Gate intramolecular ligation. Biotechnol. Bioeng. 2022.11.21.517425 (2022). doi:10.1101/2022.11.21.517425

39. Stapleton, J. A. et al. Haplotype-phased synthetic long reads from short-read sequencing. PLoS One 11, 1–20 (2016).

40. Tibshirani, R. Regression Shrinkage and Selection Via the Lasso. J. R. Stat. Soc. Ser. B 58, 267–288 (1996).

41. Hoerl, A. E. & Kennard, R. W. American Society for Quality Ridge Regression : Biased Estimation for Nonorthogonal Problems American Society for Quality Stable URL : http://www.jstor.org/stable/1267351 Linked references are available on JSTOR for this article : Ridge Regression : Biase. 12, 55–67 (1970).

42. Yuan, M. et al. Structural basis of a shared antibody response to SARS-CoV-2. Science (80-.). 369, 1119–1123 (2020).

43. Stadlbauer, D. et al. Broadly protective human antibodies that target the active site of influenza virus neuraminidase. Science (80-.). 366, 499–504 (2019).

44. Guthmiller, J. J. et al. Broadly neutralizing antibodies target a hemagglutinin anchor epitope. Nature (2021). doi:10.1038/s41586-021-04356-8

45. Throsby, M. et al. Heterosubtypic Neutralizing Monoclonal Antibodies Cross-Protective against H5N1 and H1N1 Recovered from Human IgM+ Memory B Cells. PLoS One 3, e3942 (2008).

46. Liu, L. et al. Potent neutralizing antibodies against multiple epitopes on SARS-CoV-2 spike. Nature 584, 450–456 (2020).

47. Chen, F. et al. VH 1-69 antiviral broadly neutralizing antibodies: genetics, structures, and relevance to rational vaccine design Fang. Curr Opin Virol 149–159 (2019). doi:10.1016/j.coviro.2019.02.004.V

48. Fleishman, S. J. et al. of Influenza Hemagglutinin. Science (80-.). 979, 816–822 (2011).

49. Ekiert, D. C. et al. Antibody recognition of a highly conserved influenza virus epitope : implications for universal prevention and therapy. Science (80-.). 324, 246–251 (2009).

50. Lingwood, D. et al. Structural and genetic basis for development of broadly neutralizing influenza antibodies. Nature 489, 566–570 (2012).

51. Poelwijk, F. J., Socolich, M. & Ranganathan, R. Learning the pattern of epistasis linking genotype and phenotype in a protein. Nat. Commun. 10, 1–11 (2019).

52. Vassallo, C. N., Doering, C. R., Littlehale, M. L., Teodoro, G. I. C. & Laub, M. T. A functional selection reveals previously undetected anti-phage defence systems in the E. coli pangenome. Nat. Microbiol. 7, 1568–1579 (2022).

53. Park, Y., Metzger, B. P.H. & Thornton J. W. The simplicity of protein sequence-function relationships. bioRxiv (2023). 10.1101/2023.09.02.556057

54. Hsu, C., Nisonoff, H., Fannjiang, C. & Listgarten, J. Learning protein fitness models from evolutionary and assay-labeled data. Nat. Biotechnol. 40, 1114–1122 (2022).

55. Smith, M. D., Case, M. A., Makowski, E. K. & Peter, M. Position-Specific Enrichment Ratio Matrix scores predict antibody variant properties from deep sequencing data. 1–11 (2023). doi:10.1093/bioinformatics/xxxxx

56. Ding, D. et al. Protein design using structure-based residue preferences. BioRxiv 2022.10.31.514613 (2023).

57. Wittrup, K. D., Tidor, B., Hackel, B. J. & Sarkar, C. A. Quantitative fundamentals of molecular and cellular bioengineering. (Mit Press, 2020).

58. Klesmith, J. R., Bacik, J.-P., Wrenbeck, E. E., Michalczyk, R. & Whitehead, T. A. Trade-offs between enzyme fitness and solubility illuminated by deep mutational scanning. Proc. Natl. Acad. Sci. 114, 2265–2270 (2017).

59. Engler, C. & Marillonnet, S. Golden Gate Cloning - DNA Cloning and Assembly Methods. 1116, 119–131 (2014).

60. Wrenbeck, E. E. et al. Plasmid-based one-pot saturation mutagenesis. Nat. Methods 13, 928–930 (2016).

61. Bloom, J. D. An experimentally determined evolutionary model dramatically improves phylogenetic fit. Mol. Biol. Evol. 31, 1956–1978 (2014).

62. Medina-Cucurella, A. V et al. User-defined single pot mutagenesis using unamplified oligo pools. Protein Eng. Des. Sel. 32, 41–45 (2019).

63. Ye, J., Ma, N., Madden, T. L. & Ostell, J. M. IgBLAST: an immunoglobulin variable domain sequence analysis tool. Nucleic Acids Res. 41, 34–40 (2013).

64. Medina-Cucurella, A. V & Whitehead, T. A. Characterizing Protein-Protein Interactions Using Deep Sequencing Coupled to Yeast Surface Display. Methods Mol. Biol. 1764, 101–121 (2018).

65. Chao, G. et al. Isolating and engineering human antibodies using yeast surface display. Nat. Protoc. 1, 755–768 (2006).

66. Banach, B. B. et al. Highly protective antimalarial antibodies via precision library generation and yeast display screening. J. Exp. Med. 219 (8): e20220323. (2022)

67. Kowalsky, C. A. et al. High-resolution sequence-function mapping of full-length proteins. PLoS One 10, 1–23 (2015).

