## Supporting Information for "An integrated technology for quantitative wide mutational scanning of human antibody Fab libraries"

***This PDF file contains:***
***Supplemental Note 1:* Inferring biophysical parameters from sequencing data**
***Extended Materials and Methods***
***Extended Data Figures. S1 to S10***

**SUPPLEMENTAL NOTE 1:**

#### 1. INFERRING ANTIBODY-ANTIGEN BIOPHYSICAL PARAMETERS FROM SEQUENCING DATA: EXECUTIVE SUMMARY

In deep mutational scanning, a population of mutational variants of a protein is passed through a selection or screen; this screen changes the underlying frequencies of each of these variants. Deep sequencing is used to count each variant in the population, which is used to infer the frequency of each variant in the population in a reference population and after the screen. This frequency change is converted to some score that, ideally, relates to the functional properties of the variant. This technical note describes our framework for inferring, from this processed sequence data, both dissociation constants and maximum fluorescence for antibody variants encoded in yeast displayed protein libraries screened by fluorescence activated cell sorting (FACS).

In FACS, populations are screened by collecting cells with fluorescence above a certain gating fluorescent threshold, or between two fluorescent gating thresholds; cells sorted according to these fluorescent gates are said to be sorted into bins. A clonal population of cells will exhibit a mean fluorescence with a certain variance according to cell size, surface density of displayed proteins, or other factors. Thus, only a fraction of cells for each variant will exceed the fluorescence threshold needed for collection into a given bin. Furthermore, if the fraction of cells that are sorted into a bin is known, one can infer the likely mean fluorescence for a given variant at that labeling concentration. Finally, as described in further detail below, sequencing data and other experimental observables can be used to infer the fraction of cells collected by the gating strategy and thus the mean fluorescence of a variant for a given labeling concentration. Some of the descriptions below come in part from Kowalsky et al[1], and Kowalsky et al[2].

We seek to infer variant-specific dissociation constants ($K_{d,i}$) using, for example, the Hill equation below:

$$\underline{F_i} = \left(F_{max,i} - F_{min}\right)\frac{[L]}{K_{d,i}+[L]} + F_{min} \qquad (1)$$

Here is the mean fluorescence of cells displaying variant i at a given labeling concentration [L], $F_{max,i}$ is the maximum fluorescence for the variant i, and $F_{min}$ is cellular autofluorescence.

Intuitively, if we can infer the mean fluorescence at different labeling concentrations ([L]), we can reconstruct isothermal titrations for each variant i (e.g. $\underline{F_i}$ vs. [L]) to find a best fit $K_{d,i}$ and $F_{max,i}$ using non-linear regression. An example from barcode ATGCACACATTTAAAGCTGT corresponding to variant 4A8 M59I is shown below in **Fig Note S1**.

We can approach this inference problem by regression, as it allows for the quality of the model fit to the data using the chi squared metric while also giving robust methods for confidence interval testing. As will be shown, we can also use maximum likelihood estimation in a quantitatively identical way. However, we cannot regress on the reconstructed mean fluorescence, as error is not distributed uniformly in both directions. Instead, we regress on the vector of *probabilities* of sorting into each bin.

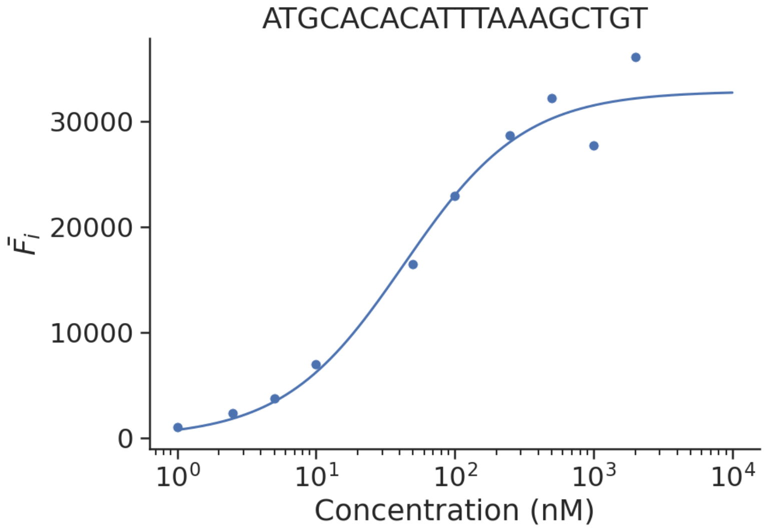

**Figure Note S1:** *Fluorescence reconstruction for barcode ATGCACACATTTAAAGCTGT at 10 labeling concentrations.*

In summary, we label our population of antibody variants at different antigen concentrations and use FACS to sort these antibody variants into different bins. Sequencing of these populations allows us to reconstruct the likely mean fluorescence for a given labeling concentration by inferring the fraction of each variant that is present in a binned population. Summing over all labeling concentrations allows us to find the most likely parameter value for dissociation constants, the confidence interval associated with that parameter estimation, and the quality of the fit using weighted nonlinear regression.

#### 2. WHAT IS THE PROBABILITY OF A CELL COLLECTED ABOVE A CERTAIN FLUORESCENT THRESHOLD?

Let's call this fluorescence threshold a gating fluorescence ($F_g$) and ask for the probability that a given clone i exhibiting a mean fluorescence intensity ($\underline{F_i}$) will be captured by this gate. A graph of this relationship is below:

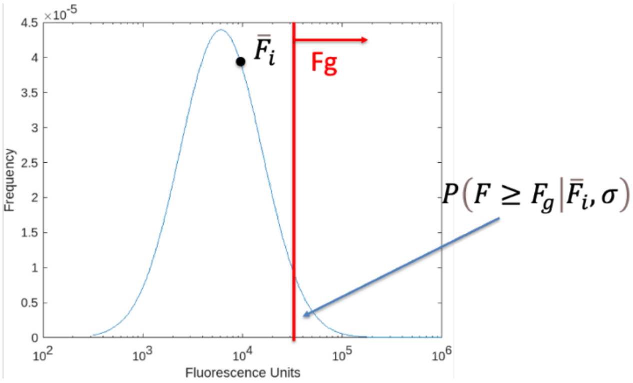

***Figure NoteS2:*** *A theoretical frequency histogram for yeast cells isogenically expressing a clone i.*

Since fluorescence measurements of clonal population of displaying cells are log-normally distributed in flow cytometry[3,4], the probability can be calculated by regular statistical calculations:

$$P(\underline{F_i}, \sigma) = \frac{1}{2} - \frac{1}{2} erf\left(\frac{lnF_g - ln\underline{F}_i + \frac{1}{2}\sigma^2}{\sigma\sqrt{2}}\right) \tag{2}$$

The other variable, $\sigma$, represents the natural log of the standard deviation of the log-normal distribution from a clonal population of cells. 'erf' is the error function used to numerically integrate a Gaussian probability distribution. Equation (2) is the fundamental equation that allows us to apply statistical calculations to derive dissociation constants.

#### 3. WE CAN FIND THE PROBABILITY OF A CELL COLLECTED BETWEEN TWO FLUORESCENT THRESHOLDS.

Assume we have a square gate set up with the lower bound some $F_{g2}$ and the upper bound $F_g$. Keeping the same definitions as above, we can rewrite a similar equation as (2) for the probability above $F_{g2}$:

$$P(\underline{F_i}, \sigma) = \frac{1}{2} - \frac{1}{2} erf\left(\frac{lnF_{g2} - ln\underline{F}_i + \frac{1}{2}\sigma^2}{\sigma\sqrt{2}}\right) \tag{3}$$

Writing the probability of that cell landing between the two gates becomes:

$$P(\underline{F_i}, \sigma) = P(\underline{F_i}, \sigma) - P(\underline{F_i}, \sigma) \tag{4}$$

$$P(\underline{F_i}, \sigma) = \frac{1}{2} erf\left(\frac{lnF_g - ln\underline{F}_i + \frac{1}{2}\sigma^2}{\sigma\sqrt{2}}\right) - \frac{1}{2} erf\left(\frac{lnF_{g2} - ln\underline{F}_i + \frac{1}{2}\sigma^2}{\sigma\sqrt{2}}\right) \tag{5}$$

Thus, the probability $p_{ijk}$ that a given cell displaying variant i can be captured in bin j at labeling concentration k is given by the following expression:

$$p_{ijk} = P(\underline{F_i}, \sigma) = \frac{1}{2} erf\left(\frac{lnF_{gjk} - ln\underline{F_i} + \frac{1}{2}\sigma^2}{\sigma\sqrt{2}}\right) - \frac{1}{2} erf\left(\frac{lnF_{g2jk} - ln\underline{F_i} + \frac{1}{2}\sigma^2}{\sigma\sqrt{2}}\right) \quad (6)$$

Note that if we have two bins with a shared boundary, we can write the *sum of the two probability distributions* as:

$$p_{ijk} + p_{ij+1k} = P(\underline{F_i}, \sigma) + P(\underline{F_i}, \sigma) = \frac{1}{2} - \frac{1}{2} erf\left(\frac{lnF_{g2} - ln\underline{F_i} + \frac{1}{2}\sigma^2}{\sigma\sqrt{2}}\right) \quad (7)$$

#### 4. WHAT DO THESE PROBABILITIES LOOK LIKE IN PRACTICE?

For a typical monovalent binding experiment, one labels yeast cells displaying a binding protein with a fluorescently conjugated ligand. We have found that $\sigma$ for phycoerythrin (SAPE) labeled populations range from 0.9-1.05 [2]. Let's assume a $\sigma = 1.02$ for this example. We find that many protein-ligand interactions we consider in lab to be well fit by a Hill equation with no cooperativity:

$$\underline{F_i} = (F_{max,i} - F_{min}) \frac{[L]_o}{K_{d,i} + [L]_o} + F_{min} \quad (1)$$

For the phycoerythrin (SAPE) labeled populations we usually consider, a typical value of $F_{min}$ representing cell autofluorescence is 350 MFI in our experimental set-up using a Sony SH800 cell sorted with a 488 nm laser and compensation for fluorescein. The two protein-specific terms are the max fluorescence ($F_{max,i}$) and the dissociation constant $K_{d,i}$ for the interaction. These will be variant-specific. For reasonably expressed and well-behaved proteins our $F_{max,i}$ is typically in the 50,000 MFI range. Let's nondimensionalize the ligand concentration so we can remove one variable.

$$\underline{F_i} = (F_{max,i} - F_{min}) \frac{\frac{[L]}{K_d}}{1 + \frac{[L]}{K_d}} + F_{min} \quad (8)$$

**Table NoteS1** reports the resulting probability lookup table:

| $\frac{[L]}{K_d}$ | $\underline{F_i}$ | $p_{ijk}$ (Fg = 2000) | $p_{ijk}$ (Fg = 5000) | $p_{ijk}$ (Fg = 10000) | $p_{ijk}$ (Fg = 25000) |
|---|---|---|---|---|---|
| 0 | 350 | .01 | <.001 | <.001 | <.001 |
| 0.1 | 4860 | .64 | .30 | .11 | .02 |
| 0.2 | 8625 | .82 | .51 | .26 | .06 |
| 0.3 | 11800 | .89 | .63 | .36 | .11 |
| 0.4 | 14500 | .92 | .70 | .44 | .15 |
| 0.5 | 16900 | .94 | .75 | .50 | .19 |
| 1 | 25200 | .97 | .86 | .65 | .31 |
| 5 | 41725 | >.99 | .94 | .81 | .50 |

This table shows that the large standard deviation resulting from square gating gives useful probabilities at many different gating fluorescence values representing different dissociation constants and/or max fluorescence values.

#### 5. WE CAN INFER THE PROBABILITY USING FREQUENCY DATA

Our observable for deep sequencing experiments is a set of read counts for variant **i** in each bin **j** and for each labeling concentration **k** (let's call these read counts $r_{ijk}$). Additionally, we have the **r**eference read counts we can observe for variant **i** (let's call this $r_{ir}$). We can directly convert observables to probabilities of sorting into a given bin j by comparing these read counts to those from the reference population. The reference population is critically important given that the comparison is the probability of being captured by a gate relative to the condition of no gate. Therefore, your reference population must be identical except for the fluorescence gate you sort at.

We write the probability as the number of cells of variant i collected in the j$^{th}$ bin and k$^{th}$ labeling concentration ($x_{ijk}$) relative to the number of cells of variant i in the reference population ($x_{ir}$):

$$p_{ijk} = \frac{x_{ijk}}{x_{ir}} \tag{9}$$

The frequency of variant i ($f_{ijk}$, $f_{ir}$) is just the number of counts observed divided by all counts, so we can write:

$$p_{ijk} = \emptyset \frac{\frac{r_{ijk}}{\sum_i r_{ijk}}}{\frac{r_{ir}}{\sum_i r_{ir}}} \tag{10}$$

Here $\emptyset$ is the total fraction of cells collected in the sorting bin relative to the reference population, and the frequency of each variant has been converted to experimental observables derived from deep sequencing. Equation (10) is the second fundamental equation because it states that the probability $p_{ijk}$ (set by $F_{max,i}$ and $K_{d,i}$) we observe for a given labeling concentration k and bin j are a function of the observables from the deep sequencing experiment.

#### 6. SOURCES OF NOISE IN RECONSTRUCTING FLUORESCENCE FROM EXPERIMENTAL OBSERVABLES

A major challenge for sequence-function reconstruction experiments comes from determining the appropriate confidence level set for each experimental measurement. This is important as low and high values of $p_{ijk}$ give large uncertainties in the measurement of $\underline{F}_{ik}$ (see **Fig Note S3** below).

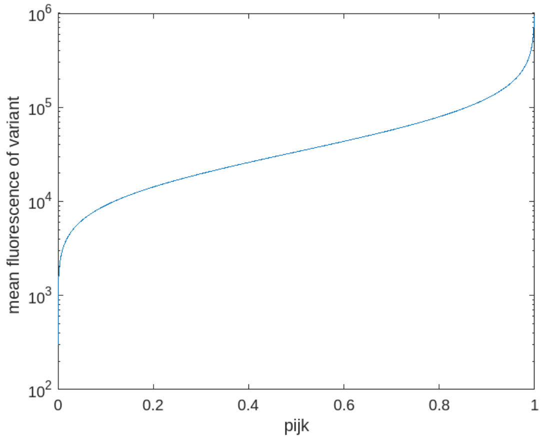

**Figure Note S3:** *Mean fluorescence of variant as a function of* $p_{ijk}$ *- the probability of sorting into a bin above some* $F_g$ *= 30,000 MFI. Low and high observed probabilities result in large, one-tailed uncertainties in the value of the mean fluorescence.*

For parameter inference it is important to identify and quantify sources of noise in the fluorescence reconstruction. Intrinsic noise comes in the act of sorting discrete cells, preparing amplicons from yeast cells by PCR, and sequencing discrete nucleic sequences. Extrinsic noise results from the efficiency of the overall process of cell sorting and recovery.

We have previously shown[5] that deep sequencing read counts from FACS data can be evaluated according to Poisson probability distributions. We have previously determined the propagation of errors for Poisson noise in the counts of the reference and selected populations[6]. Although this is an underestimate of error because population bottlenecks occur during cell sorting, outgrowth, and amplicon prep leading to overdispersion, we find empirically that Poisson noise is a reasonable approximation for well-designed experiments. By propagation of errors, we can determine the variance associated with the value of $p_{ijk}$:

$$\sigma^2_{pijk,intrinsic} = \sigma^2_{rijk}\frac{\partial pijk}{\partial rijk}^2 + \sigma^2_{rir}\frac{\partial pijk}{\partial rir}^2 \tag{11}$$

We can approximate the Poisson noise as

$$\sigma^2_{rijk} = r_{ijk};\ \sigma^2_{rir} = r_{ir} \tag{12}$$

$$\sigma^2_{pijk,intrinsic} = r_{ijk}\left(\emptyset\frac{\frac{1}{\Sigma_i \quad r_{ijk}}}{\frac{r_{ir}}{\Sigma_i \quad r_{ir}}}\right)^2 + x_{ir}\left(-\emptyset\frac{\frac{r_{ijk}}{\Sigma_i \quad r_{ijk}}}{\frac{{r_{ir}}^2}{\Sigma_i \quad r_{ir}}}\right)^2 \tag{13}$$

$$\sigma^2_{pijk,intrinsic} = \frac{{p_{ijk}}^2}{r_{ijk}} + \frac{{p_{ijk}}^2}{r_{ir}} \tag{14}$$

The other source of error is extrinsic relating to error rate in the sorting itself – what is the probability of a mis-sorting event? This appears to be 2% in our experimental set-up, but we expect that this error rate may vary.

$$\sigma^2_{pijk,extrinsic} = (0.02)^2 \tag{15}$$

In the experiments presented in Figures 4 and 5 of the main text, we included Fab nonbinders to measure this error directly from the sequencing data. For these experiments, these values were observed to be 0.014, close to the values used in the initial experiment.

Taken together by propagation of error, we end up with the following result for the uncertainty associated with probability:

$$\sigma_{pijk} = \sqrt{\sigma^2_{pijk,extrinsic} + p_{ijk}{}^2\left(\frac{1}{r_{ijk}} + \frac{1}{r_{ir}}\right)} \tag{16}$$

#### 7. PARAMETER ESTIMATION USING MAXIMUM LIKELIHOOD ESTIMATION

The log likelihood framework states that the parameter set most likely to fit a given set of data occurs with maximization of the summation of the log probabilities of each experimental measurement:

$$LL_i(K_{d,i}, F_{max,i}) = \left(\sum_{jk} \quad logP\left(Model_{ijk}\right)\right) \tag{17}$$

Here, $Model_{ijk}$ is the model probability (given parameters $K_{d,i}, F_{max,i}$) of a variant i being sorted into bin j at labeling concentration k. We must assume some probability distribution – given the sources of noise and the fact that reference and sorted counts are typically >10, a Gaussian probability distribution is justifiable here. Expanding terms, we can write:

$$LL_i(K_{d,i}, F_{max,i}) = \left(\sum_{jk} \quad logGaussianPDF\left(Model_{ijk}\right)\right) \tag{18}$$

Expanding the Gaussian probability distribution and removing constant terms, we arrive at:

$$LL_i(K_{d,i}, F_{max,i}) = \sum_{jk} \quad -\frac{1}{2}\left(\frac{p_{ijk} - Model_{ijk}}{\sigma_{ijk}}\right)^2 \tag{19}$$

Note that maximizing this expression is equivalent to minimizing the weighted sum of square errors or the chi squared metric. The algorithm changes the probabilities of $Model_{ijk}$ by changing parameters in the Hill function, and we use off-the-shelf optimization software to find the minimization of the function.

$$-LL_i(K_{d,i}, F_{max,i}) = \sum_{jk} \quad \left(\frac{p_{ijk} - Model_{ijk}}{\sigma_{ijk}}\right)^2 \tag{20}$$

#### 8. CONFIDENCE INTERVALS USING MAXIMUM LIKELIHOOD ESTIMATION

Using a MLE framework that minimizes the chi squared metric ($\chi^2_{min}$) results in the simplification of confidence interval measurements. As such, we follow standard approaches[7,8] for determining 95% confidence intervals using the critical value of the F distribution statistic ($F_{0.05}$) using the following equation:

$$\frac{\chi^2}{\chi^2_{min}} \leq \frac{n-2}{n-1}\left(1 + \frac{n}{n-1} F_{0.05}\,(n-1, n)\right) \qquad (21)$$

Where n is the number of experimental data points (here, the number of bins used for MLE), and $\chi^2$ is the chi squared metric for given parameter values of $K_{di}$ and $F_{maxi}$.

### *Extended Materials and Methods*

#### Plasmids

All plasmids were constructed using either NEBuilder HiFi DNA Assembly Master Mix (New England Biolabs) for Gibson assembly[9], by Golden Gate assembly[10,11], using a Q5 Site-Directed Mutagenesis Kit (New England Biolabs), or by nicking mutagenesis[12,13]. Synthetic DNA was ordered either as gBlocks or eBlocks (IDT). A complete list of plasmids, libraries, gene blocks, and primers are located **Supporting File 1**.

pMBK046, the old 4A8 Fab YSD vector, was constructed by Golden Gate assembly with plasmids pMBK008 and pMBK027 and gene blocks 7 & 8. pMBK047-pMBK228, the 4A8 Fab YSD library plasmids, were generated by combinatorial nicking mutagenesis[12] with pMBK046 as the template and primers 142 and 328-339 and isolated by Sanger sequencing (Genewiz) of individual colonies.

The mini-mutagenesis shuttle vectors for the 4A8/CC12.1/COV2-2489 library: pMBK231, pMBK233, pMBK234, pMBK235, pMBK236, pMBK237, pMBK248 were generated by golden gate assembly of the antibodies corresponding VH and VL gene fragments with pBMP103-UMI and pBDP and were sequence confirmed by Oxford nanopore sequencing (Plasmidsaurus).

The isogenic yeast display Fab plasmids were constructed by yeast homologous recombination. The sequence verified shuttle vector(s) (pMBK231 – pMBK248, pMBK317 - pMBK318, pMBK341 - pMBK342) and the yeast display plasmids pBMP103 for kappa antibodies and pMBK275 for lambda antibodies were separately digested with NotI-HF and bands corresponding to the antibody Fab or yeast vector backbone were fractionated on an agarose gel and extracted using Macherey-Nagel NucleoSpin® Gel and PCR Clean-up kit (740609.50). The purified DNA was mixed in a 2:1 molar ratio of Fab insert to yeast display backbone and co-transformed into chemically competent EBY100.

The Fab shuttle vectors with kanamycin (pBMP101 and pMBK272) and chloramphenicol (pMMP_kappa and pMMP_lambda) antibiotic resistance genes were constructed using either gBlocks or eBlocks (IDT) and Gibson Assembly using NEBuilder HiFi DNA Assembly Master Mix.

The lambda mRFP yeast surface display plasmid, pYSD_lambda_mRFP, was constructed by first digesting pYSD_kappa_mRFP with PacI and NotI-HF to remove the kappa light chain segment and next the lambda light chain e-block was cloned in by Gibson Assembly using NEBuilder HiFi DNA Assembly Master Mix.

#### Construction of Fab libraries

To generate libraries L001, L002, L003, L024, and L029, combinatorial mutagenesis was performed exactly as previously described[12,14] using the mini-mutagenesis plasmids pMBK234, pMBK235, pMBK236, pMBK237, pMBK233, pMBK248, pMBK317, and pMBK318, as the parental plasmid DNA templates and the mutagenic oligos 137, 139, 142, 145, 146, 153, 332-339, 347-356, 375-376, and 456-459, (IDT) containing degenerate codons that encode either for the mature antibody residue or the UCA residue.

To construct libraries OMIL004, OMIL006, OMIL011, and OMIL0012 for 319-345, 222-1C06, 1G01, and 1G04, targeted site-saturation mutagenesis was performed by the method of Bloom[15]. Template linear PCR products were made using primers OMI1009 and OMI1011 which amplify the $V_H$-BDP-$V_L$ region of plasmids OMI0014, OMI0016, OMI0020, and OMI0021. 25 µL of 2x Q5 Master Mix, 2.5 µL of 10 µM OMI1009, 2.5 µL of 10 µM OMI1011, 1.2 µL of 5 ng/µL of the template plasmid were combined with 18.8 µL of water. Each of the PCR reactions were run on a thermocycler at the following settings:

1. 98 °C for 30 s.
2. 98 °C for 10 s.
3. 72 °C for 56 s.
4. Repeat steps 2 and 3 for 24 additional cycles.

PCR products were then purified over agarose gels using a NucleoSpin® Gel and PCR Clean-up kit (Macherey-Nagel, 740609.50). These products were used as templates for the forward and reverse fragmenting reactions prepared as described[15] with the following modifications: 2x Q5 Master Mix was substituted for 2x KOD Hot Start Master Mix, OMI1009 was used as the outer forward primer, OMI0011 was used as the outer reverse primer, and mutagenic oligos tiling the $V_H$ and $V_L$ CDRs were used for the forward and reverse pools. The reactions were run for 10 cycles of the above thermal cycler program. The products from this reaction were used as templates for the joining reaction. The joining reaction was prepared as described in Bloom with the same modifications from the fragmenting reactions. Additionally, the joining reaction volume was scaled from 30 µL to 50 µL to increase product yield. The joining reaction was cycled using the above program for 20 total cycles. The products from this reaction were gel purified over agarose gels using a NucleoSpin® Gel and PCR Clean-up kit (Macherey-Nagel, 740609.50).

L030, the UCA_2-17 forward trajectory library was generated following oligo pool mutagenesis[16] with a fast anneal step performed with the oligo pool listed in **Supporting File 1**. The molar ratio of template DNA to oligonucleotides was adjusted to 10:1.

1 μg of plasmid libraries in a reaction volume of 20 μL in rCutSmart buffer were separately digested with 20 units of NotI-HF (NEB) for 1 hour at 37 °C. In parallel, 2-5 μg of pBMP103-UMI or pMBK275-UMI library in a reaction volume of 20 μL in rCutSmart buffer was also digested with 20 units of NotI-HF for 1 hour at 37 °C. The digested DNA was fractionated on a 1 (w/v) % agarose gel. Bands corresponding to the $V_H$-BDP-$V_L$ region of the antibodies (1.8 kB) and the yeast surface display vector backbone for the UMI library (6.4 kB) were extracted using Macherey-Nagel's NucleoSpin® Gel and PCR Clean-up kit (Macherey-Nagel, 740609.50).

The yeast surface display and barcoded mutagenic antibody library (L006 4A8/CC12.1/COV2-2489) was generated using Gibson Assembly with the gel extracted components in a 2:1 molar ratio of antibody library insert to pBMP103-UMI yeast surface display vector. Each of the antibody combinatorial libraries contained 64 variants and were mixed in an equimolar amount for the Gibson Assembly reaction[9] using the NEBuilder HiFi DNA Assembly master mix following the manufacturer's protocol. A column clean-up was performed to remove residual buffer and enzymes and approx. 25% of the cleaned reaction product was transformed into homemade chemically competent *E. coli* Mach1. The next day, 11,000 transformants were observed from the transformation dilution plate and the entire library was harvested and miniprepped.

The barcoded yeast surface display libraries for the S1/HA mixed antigen sort were generated by Gibson assembly of the gel extracted components in a 2:1 molar ratio of antibody library insert to yeast surface display vector for the kappa and lambda antibody libraries separately. L024 and L029 were assembled with L018 library and L030, OMIL004, OMIL006, OMIL011, and OMIL0012 were assembled with pBMP103-UMI library in a total reaction volume of 20 μL. Both Gibson Assembly reactions were incubated for 4 hours at 50 °C and the reaction products were cleaned and concentrated to 6 μL with a Monarch DNA & PCR Cleanup Kit (NEB). The entire 6 μL product was transformed via electroporation into TransforMAX cells (Lucigen, EC300110) and incubated at 37 °C overnight. A dilution plate was used to assess the transformation efficiencies and the transformants were bottlenecked to 2,000 lambda variants and approximately 8,000 kappa variants. The bottlenecked libraries were grown up in 50mL SOB + kanamycin overnight at 37 °C. The next day the libraries were miniprepped and pooled together and 6 different 4A8 barcoded plasmids were also spiked into the library pool. 5 μg of plasmid DNA was transformed into chemically competent EBY100 in parallel reactions for the biological replicate libraries.

#### Barcode-variant pairing

Barcodes were paired with $V_H$ and $V_L$ variants through Oxford nanopore sequencing or by short read sequencing of short read amplicons prepared by intramolecular ligation of barcode UMI in proximity to the CDR3 of either the $V_H$ or $V_L$. Oxford nanopore sequencing (Plasmidsaurus) was performed on individual plasmids. Short-read amplicons were sequenced on an Illumina MiSeq with 2x250 bp paired end reads (Rush University Sequencing Core).

The optimized intramolecular ligation procedures were performed in a reaction volume of 100 μL with 1 μg of plasmid library, 400 U of T4 DNA ligase (1 μL of 400 U/μL), and 1X T4 DNA ligase buffer (NEB; catalog # B0202S). Also added to the reaction were either 30 U of BbsI (3 µL of 10 U/µL) (NEB; catalog # R0539L) - for the intramolecular ligation of barcode to VL - or 30 U of PaqCI (3 µL of 10 U/µL) plus 3 µL PaqCI activator (diluted from 20 µM stock 1:4 in 1X T4 DNA ligase buffer) (NEB; catalog # R0745L) - for ligation of barcode to VH. The reaction was subjected to 60 cycles of 37 °C for one minute followed by 16 °C for one minute, and then a final incubation of 37 °C for 5 minutes. Exonuclease III was added to each reaction (1 µL of a 1:10 dilution made in 1X rCutSmart buffer from 100 U/µL stock) (NEB; catalog # M0206L) followed by incubation at 37 °C for 30 minutes. We performed electrophoresis on the reactions on 1% (w/v) agarose gels in 1X TAE and gel extracted (Macherey-Nagel, catalog # 740609.50) the bands corresponding to intramolecular ligated products for the $V_H$-barcode and $V_L$-barcode pairing.

Amplicons were prepared by first performing a PCR using primers 428 - 431 to amplify the barcode and gene sequence (UMI-$V_L$: 428 & 430; UMI-$V_H$: 429 & 431) and append Illumina TruSeq small RNA compatible sequences. 10 ng of gel extracted input DNA was amplified in a 25 µL total reaction with Phusion High-Fidelity Polymerase with reaction components following the manufacturers recommended protocol. The PCR thermocycler program progressed for 12 cycles. After the first PCR was completed, a shrimp alkaline phosphatase (rSAP) clean-up step was performed according to the manufacturer's instructions on 10 µL of PCR product. 1 µL of rSAP cleaned DNA was used as the input for the seconod PCR, which amplifies the amplicon further and appends unique 6-bp TruSeq small RNA barcodes and Illumina sequencing adapters. The second PCR was performed using Phusion High-Fidelity Polymerase with reaction components following the manufacturers recommended protocol for 14 cycles with a 25 µL total reaction volume. After the second PCR, the amplicons were cleaned with Ampure XP beads (Beckman Coulter, A63880) following the manufacturer's instructions. Each dsDNA PCR product was quantified using Quant-IT PicoGreen (Invitrogen, P11495) and pooled for deep sequencing.

For optimization of the protocol above, three individual plasmids were sequenced by Oxford nanopore. These plasmids were then mixed in pre-defined ratios and different amplicon prep conditions were applied. These differences include: the polymerase used, the number of PCR cycles, and the type of PCR clean-up between the first and second PCRs. The three polymerases used were Phusion High-Fidelity DNA polymerase (NEB, M0530), Q5 polymerase in a 2X Master Mix (NEB, M0492), and KAPA polymerase (Roche, 7958927001) and each PCR reaction components and thermocycler program were performed according to the manufacturer's instruction.

#### Amplicon Preparation and Deep Sequencing

1e6 – 4e6 sorted yeast cells from each collected population were miniprepped according to Medina-Cucurella & Whitehead[17] using Zymoprep Yeast Plasmid Miniprep kits in either individual Eppendorf tubes (D2004) or 96-well plate format (D2007) and plasmid DNA was eluted in 30 µL nuclease free water. 15 µL of eluted plasmid DNA was further purified with exonuclease I and lambda exonuclease. The UMI region of the purified DNA was amplified using 25 PCR cycles with Illumina TruSeq small RNA primers following Kowalsky et al. 'Method B'[1] using Phusion High-Fidelity DNA polymerase in a 50 μL total reaction. 5 μL of the PCR product was size verified on a 1% (w/v) agarose gel and the remaining 45 μL was cleaned with Ampure XP beads (Beckman Coulter, A63880) following the manufacturer's instructions. Each dsDNA PCR product was quantified using Quant-IT PicoGreen (Invitrogen, P11495) and pooled in equimolar amounts for deep sequencing. The sorted UMI's were sequenced on either an Illumina MiSeq (4A8/CC12.1/COV2-2489 sort) or NovaSeq6000 (S1/HA sorts) by Rush University with single end reads.

#### Data Processing

Sequencing files were processed using the custom Python code accessible on GitHub (https://github.com/WhiteheadGroup/MAGMA-seq). The code contains three primary modules used in this work referred to as *haplotyping*, *scanning*, and *parameter estimation*. *Haplotyping* takes input sequencing files from internally ligated yeast display plasmids and creates a barcode-to-variant map. *Scanning* reads input sequencing files for sorted yeast populations for which only barcode sequences are processed, counts, and matches the barcodes to a variant specified in the previously generated barcode-to-variant map, and integrates this with sorting conditions for final output. Finally, the *parameter estimation* module performs maximum likelihood estimation (MLE) on each variant contained in the output to generate parameter estimates for $K_D$ and $F_{max}$ with 95% confidence intervals determined from reduced chi squared. See the "config" folder on Github for exact parameters used for generating each of the datasets used in this work.

For each of the sequencing processing modules (haplotyping and scanning), FASTQ files and processing parameters are entered into a config file (see README and example config files on GitHub repository). Necessary packages including Biopython, NumPy, and SciPy can be easily installed into a conda virtual environment with the included YAML file. The code is highly efficient and parallelizable (using the multiprocessing library) and can run on datasets containing millions of sequences in under an hour on our hardware (Alpine supercomputing cluster (CU Research Computing) x86_64 AMD Milan CPU with 32MB L3 Cache (utilizing 8 cores), 3.75 GB RAM/core).

#### Sequence merging and filtering

We adapted the software from Haas et al. 2021[18] for merging paired end reads at all the pertinent amplicon lengths. Sequences are then filtered based on the sequence agreement within overlap regions (see Haas et al. 2021 for algorithm details) as well as overall minimum quality scores across the full amplicon, gene, and barcode regions. Barcodes and genes are extracted from these successfully merged reads assuming a fixed location within the amplicon. Sequences are then collapsed and counted based on unique barcode and gene combinations and gene sequences are matched to a set of possible wild-type sequences based on Hamming distance. Amino acid mutations are then determined based on the chosen wild-type sequence. Variant frequencies are calculated by considering the genotypes represented in the dataset, ignoring barcodes.

A six-letter sequence motif (**CGGCGG**) occurring within the COV2-2489 antibody $V_H$ gene causes a precipitous drop in quality scores of all base calls downstream from this previously known motif[19] on the reverse read (**Figure S5**). This drop in quality score required a different haplotyping protocol for this antibody library where paired reads were not merged. This is justifiable as the mutations encoded in our library all exist on the high quality forward read, and the barcodes are located on the reverse read upstream of the quality drop. We identified reads from this antibody by matching the read to an unmutated region at the beginning of the sequence ($V_H$ positions 1-31), filtered the individual forward reads based on quality (dropping reads with overall minimum quality <Q10), and then paired the associated barcode from the reverse read based on index. The resulting data was passed into the following haplotyping step identical to the other merged reads.

#### Haplotyping: pairing a barcode barcode with Fab variant

For each barcode, we make a variant call by comparing all observed barcode-genotype pairings. We first apply a mutation filter (variants with more than 10 mutations from the assigned wild-type are removed). We then apply the count filter and a frequency filter (see config for filtering values used). Additionally, options are available for removing all silent mutations and/or all mutations not encoded in the mutagenic library. From the remaining possible pairings, we divide the observed read counts by the variant frequency and select the variant with the maximum of these values. This method appropriately weights lower observed counts of rarer variants. The raw pairing data as well as the processed barcode-to-variant map are output to CSV files for analysis.

After barcode maps have been made for $V_H$ and $V_L$ segments, we merge the two maps based on identical barcodes and output the resulting pairings as a CSV file. A final check is performed where barcodes that pair to heavy and light chains from distinct antibodies are removed from the map, resulting in a **barcode-to-variant** map.

#### Scanning

The *scanning* module matches sequenced barcodes from sorted libraries to barcodes in the map produced by the haplotyping module. It is designed to process single reads only (alternatively, forward reads from paired-end sequencing runs can be used). As described previously, barcodes are identified based on fixed location within the read. As a default, barcodes are filtered based on adherence to the template sequence of mixed bases; this option can be turned off. Each barcode is matched to an identical barcode in the barcode-to-variant map and the number of times each barcode is seen is summed. Information entered by the user in “limit.csv” including high and low bin limits (Hj and Gj, respectively), and number of cells sorted and collected in each bin (Nk and Njk, respectively) are matched with concentration and bin names to generate the CSV file needed for the next parameter estimation step. Note that the concentration and bin names in the “limit.csv” must match the identifiers from the scanning barcodes config file. For example, a bin named “top25” at concentration “5nM” should be identified in the config file as “conc5nM_bintop25”.

Scan barcode outputs information for each of the concentrations and bins analyzed as well as an overall output in two forms: (1) "_combined.csv" records the counts by barcode and (2) "_collapsed.csv" records the counts collapsed by antibody variant. Both the combined and collapsed CSV files are ready for input into the parameter estimation module assuming a proper "limit.csv" was specified. For each bin and concentration, the percentage of barcodes that matched to a variant in the map is recorded. With our conservatively filtered maps, these percentages tend to range between 50-70% read matching. Far lower percentages usually indicate low efficiency of haplotyping.

#### Parameter estimation

The parameter estimation module generates maximum likelihood estimates for $F_{max}$ and $K_D$ for each barcoded variant. Parameter estimation requires that the data be supplied in the format of the scan barcode output (see examples/scan_output.csv for example) where each row specifies a single observation of a variant labeled at a given concentration and collected at a given bin. Additionally, a few global parameters must be entered by the user. First, sigma defines the width of the lognormal distribution that represents the possible fluorescence range for a given variant. We have found that this value is somewhat independent of the variant and label concentration. In our testing, these values range from 0.90-1.02. Second, "B" represents the variant-independent cell autofluorescence, which can be determined by reading fluorescence values of Fab-expressing yeast cells without binding partner. We find that this value should fall in the range of 290-350 RFU (in PE channel) using our equipment (Sony SH800, yeast cells, 488 nm laser with compensation for PE/AlexaFluor488 fluorophores).

To achieve accurate estimates for weak binders and poorly expressed Fabs (high $K_D$ or low $F_{max}$) a few modifications to the MLE algorithm were necessary. First, manual curation was used to remove bins that had poor sequencing coverage or that had inferred probabilities that were inconsistent with the other datasets. For the data represented in Figure 2 and 3, the two bins affected were at the 25 nM labeling concentrations. The MLE algorithm first performs parameter estimation using all remaining top 25% bins. The maximum likelihood estimates were analyzed and variants with calculated $K_D$ that exceeded 1μM or $F_{max}$ that fell under 12,000 RFU were removed. For these variants, MLE was performed again by concatenating top 25% and next 25% bins into a single top 50% bin at each concentration using the joint probability estimate using equation (7). For the mixed antigen sorts represented in Figures 4 and 5, 2-7 variants were assessed by combining both bins for 25 and 50 nM labeling concentrations. Additionally, all anti-S1 probabilities $p_{ijk}$ were multiplied by 0.64 to correct for cell sorter efficiency in this experiment.

### Extended Data Figures

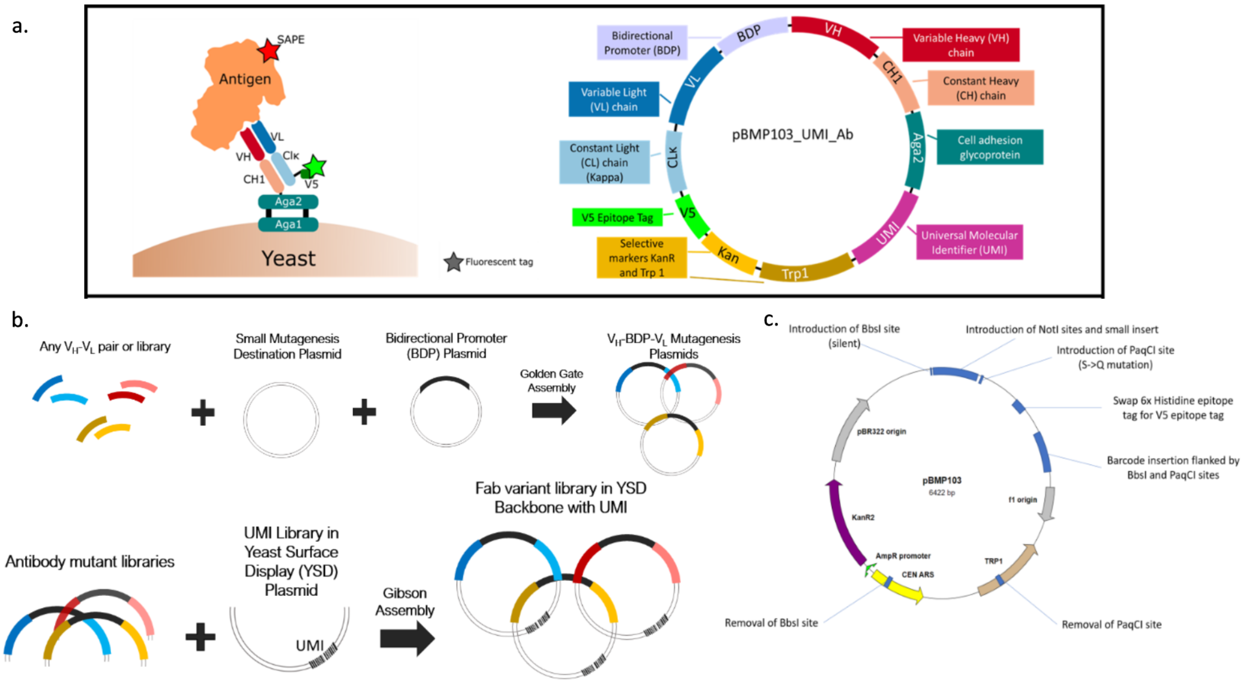

**Extended Data Figure 1 | Schematic of assembly strategy with shuttle vector and yeast display mutations. a.** Yeast display plasmid map highlighting most of the relevant genes (shown is kappa only; the lambda map is otherwise identical). The gene segments are not drawn to scale. **b.** Two-step cloning strategy for assembling barcoded Fab libraries. Along with a bidirectional promoter (BDP) plasmid, any $V_H$-$V_L$ pair or library is assembled by Golden Gate into a minimal 3.6 kB mutagenesis plasmid containing a $Cam_R$ selection marker, a high copy number ORI, and regions of homology to the CH1 and light chain sequence. There are small mutagenesis destination plasmids for both human kappa and lambda light chains. After mutagenesis, antibody mutant libraries are assembled by the method of Gibson with a yeast surface display plasmid containing a unique UMI. This final plasmid is identical to that in panel a. **c.** Summary of mutations in the yeast display vector. The major missense change is S5Q on CH1 necessary for encoding a PaqCI restriction site near the CDR H3 for short read sequencing pairing of UMI to the $V_H$ gene. We also removed the light chain 6x Histidine epitope tag and replaced it with a V5 epitope tag (GKPIPNPLLGLDST) to be able to measure binding to antigens that may be His-tagged with an anti-His conjugated fluorophore. Unique PaqCI and BbsI sites are necessary for UMI-Fab pairing by short read sequencing; antibody sequences encoding these sites are not compatible with the short read sequencing protocol.

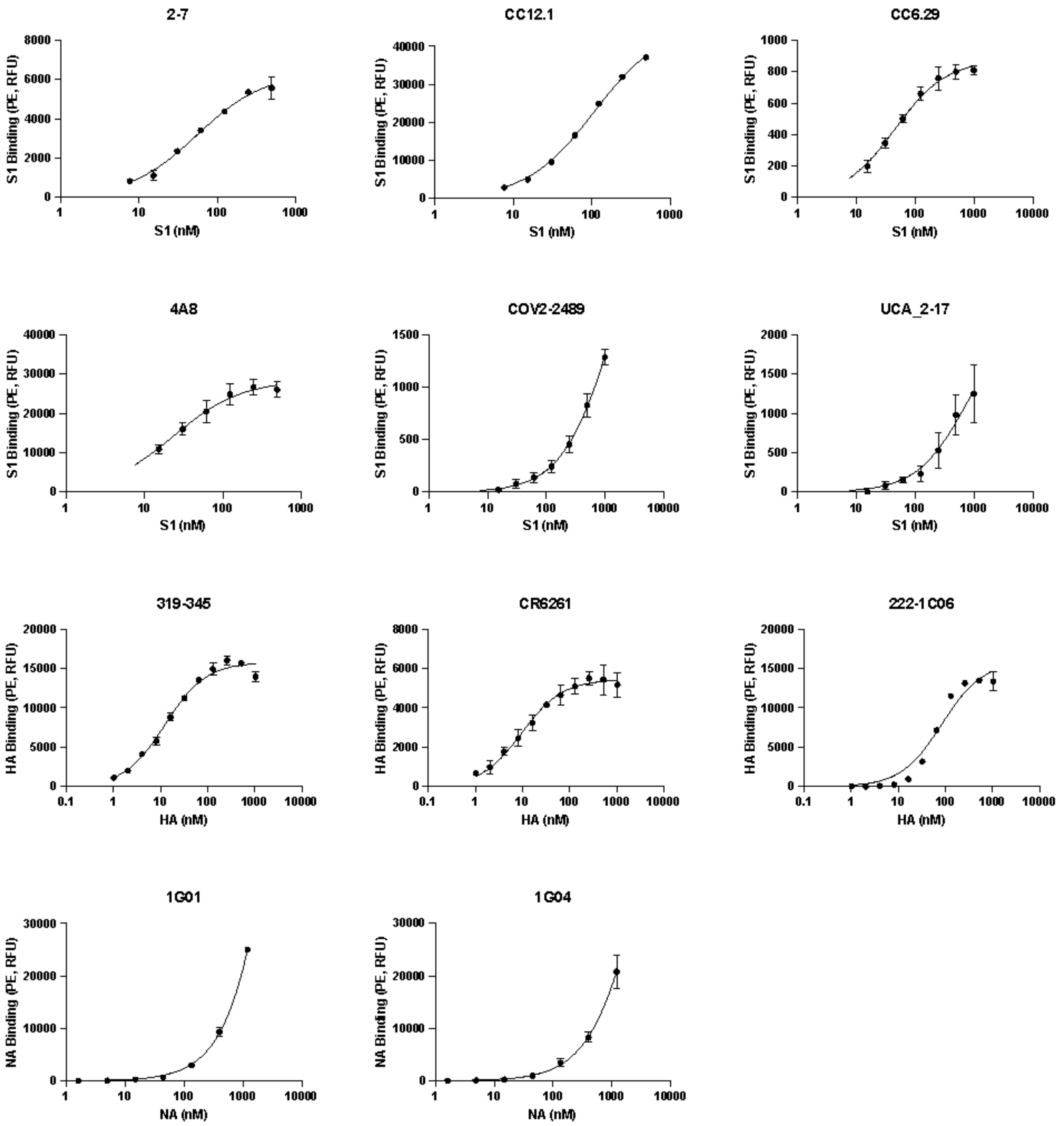

**Extended Data Figure 2 | Yeast surface display titrations.** Isogenic yeast surface titrations for antibodies reported in main text Figure 1c. Error bars represent 1 s.d. of 2 measurements. The curve fit shown is a Hill equation where the Hill coefficient is constrained to unity.

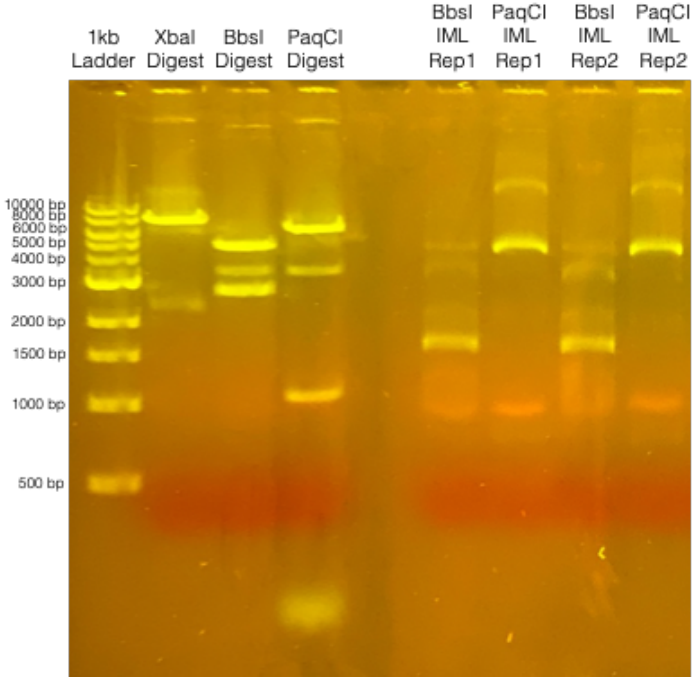

**Extended Data Figure 3 | Intramolecular Ligation Products.** Intramolecular ligation (IML) reaction products from 1 µg of barcoded 4A8/CC12.1/COV2-2489 Fab library (lanes 6-9) were separated by a 1% (w/v) agarose gel. Lanes 2-4 show individual controls reactions without ligase. The ligation product from BbsI is 1.8 kB, while that from PaqCI is 6.4 kB. In these reactions, the UMI is paired to the $V_L$ with BbsI intramolecular ligation and to the $V_H$ with PaqCI. Biological replicates (Rep1, Rep2) were performed, yielding reproducible band sizes and intensities.

| | | | True Frequencies of Correct Pairing | | | P values (T test) | | | | | | |
|---|---|---|---|---|---|---|---|---|---|---|---|---|
| Polymerase | PCR Cycles | PCR Clean-up Method | Replicate 1 | Replicate 2 | Replicate 3 | Q5, 25x Cycles, rSAP | Phusion, 12x, rSAP | KAPA, 14x, rSAP | Q5, 14x, rSAP | Q5, 14x, Column | KAPA, 14x, Column | Phusion, 12x, Column |
| Q5 | 25 | rSAP | 60% | 59% | 55% | 5.00E-01 | | | | | | |
| Phusion | 12 | rSAP | 90% | 92% | 91% | 1.24E-05 | 5.00E-01 | | | | | |
| KAPA | 14 | rSAP | 89% | 87% | 86% | 3.34E-05 | 8.99E-03 | 5.00E-01 | | | | |
| Q5 | 14 | rSAP | 82% | 83% | 81% | 6.68E-05 | 3.29E-04 | 7.13E-03 | 5.00E-01 | | | |
| Q5 | 14 | Column | 85% | 78% | 79% | 5.43E-04 | 5.16E-03 | 2.57E-02 | 2.75E-01 | 5.00E-01 | | |
| KAPA | 14 | Column | 86% | 89% | 92% | 6.74E-05 | 1.29E-01 | 2.13E-01 | 1.00E-02 | 1.98E-02 | 5.00E-01 | |
| Phusion | 12 | Column | 82% | 84% | 78% | 3.10E-04 | 3.75E-03 | 2.38E-02 | 3.53E-01 | 4.09E-01 | 1.90E-02 | 5.00E-01 |

| | | | True Frequencies of Correct Pairing | | | P values (T test) | |
|---|---|---|---|---|---|---|---|
| Polymerase | PCR Cycles | PCR Clean-up Method | Replicate 1 | Replicate 2 | Replicate 3 | Q5, 25x Cycles, rSAP | IML Control, 25x Cycles, rSAP |
| Q5 | 25 | rSAP | 60% | 59% | 55% | 5.00E-01 | |
| IML Control Q5 | 25 | rSAP | 64% | 73% | 65% | 2.10E-02 | 5.00E-01 |

**Extended Data Figure 4 | Optimization of PCR amplicon preparation for barcode-Fab haplotyping.** Three isogenic plasmids (one CC12.1 variant, two 4A8 variants) were mixed at different molar ratios and the intramolecular ligation for the $V_L$ chain and amplicon protocols were performed in triplicate (n=3). We varied the following parameters: polymerase, number of PCR cycles, and PCR clean-up method (rSAP (New England Biolabs) or column cleanup). Amplicons were deep sequenced on the same MiSeq flow cell, and observed frequencies of pairing were extracted. True frequencies of correct pairing ($freq_{true}$) were obtained for the lowest abundant variant using the following equation:

$$freq_{true} = \frac{freq_{obs} - f}{1 - f} \quad (22)$$

Here, $freq_{obs}$ is the observed frequency by deep sequencing and f is the fraction of the lowest abundant variant. As this fraction goes to zero the true frequency is identical to the observed frequency. True frequencies and p-values from paired, one-tailed t-tests are reported. We also report true frequencies and p-values of performing the intramolecular ligation separately on individual plasmids (IML control). The protocol chosen for barcode-Fab haplotyping is highlighted in green.

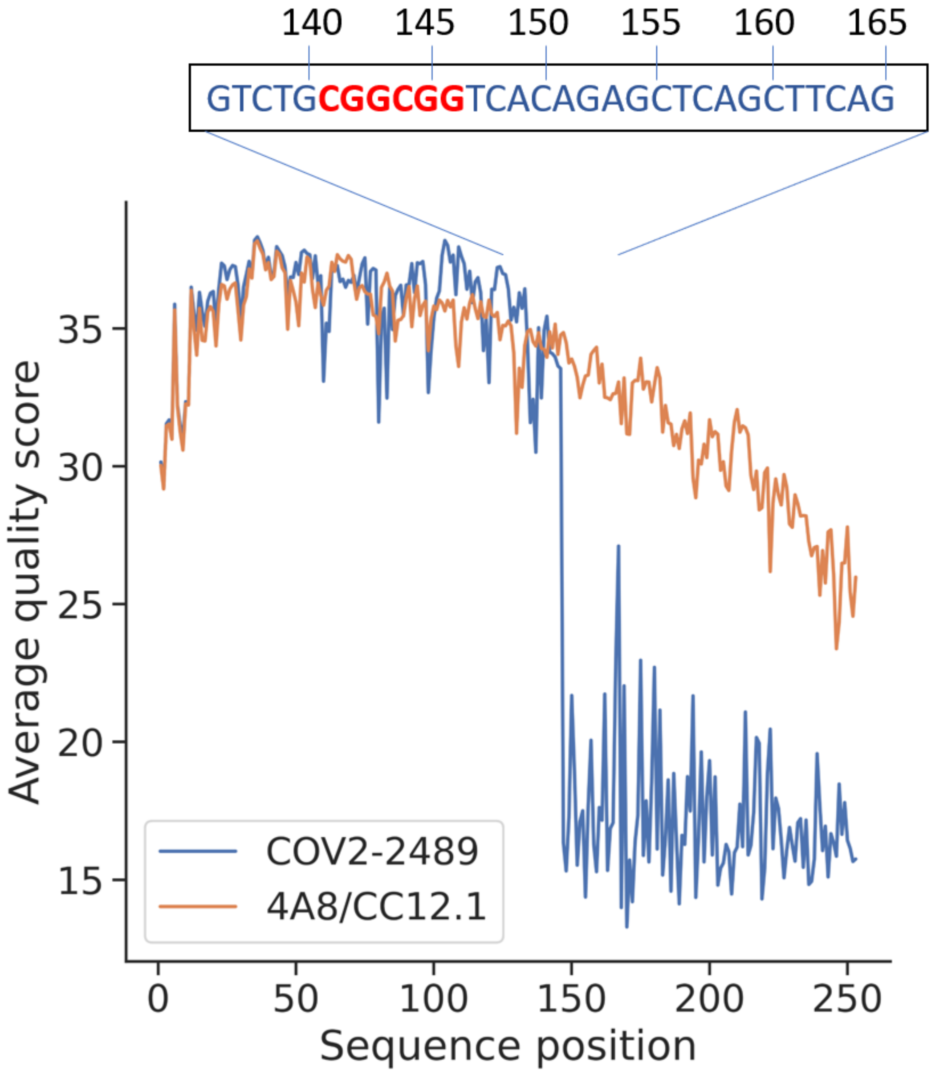

**Extended Data Figure 5 | Repeated 'CGGCGG' motif in COV2-2489 sequences causes drop in quality at nucleotide 147 on $V_H$ reverse read.** Average quality score vs. sequence position for 4A8 & CC12.1 antibodies (orange) compared to COV2-2489 (blue). The inset shows the nucleotide sequence adjacent to the drop in quality score.

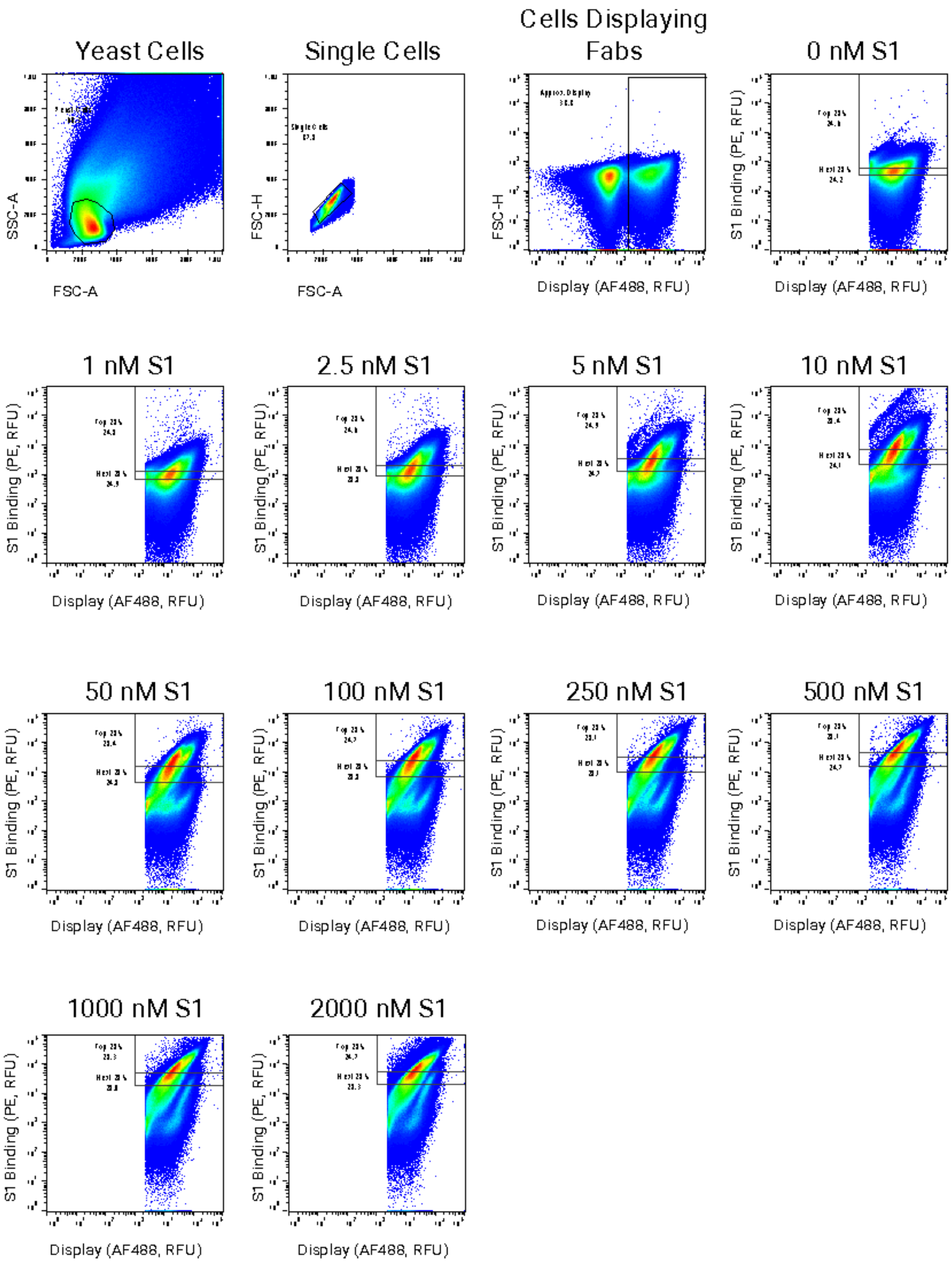

**Extended Data Figure 6 | Cytograms from first sort with S1 and 4A8/CC12.1/COV2-2489 Antibodies.** Cytograms showing sorting gates for first demonstration of the method with mixed Abs against S1. Cells were first gated for yeast cells, single cells, and cells displaying Fabs before being gated and sorted for the top 25% and next 25% bins.

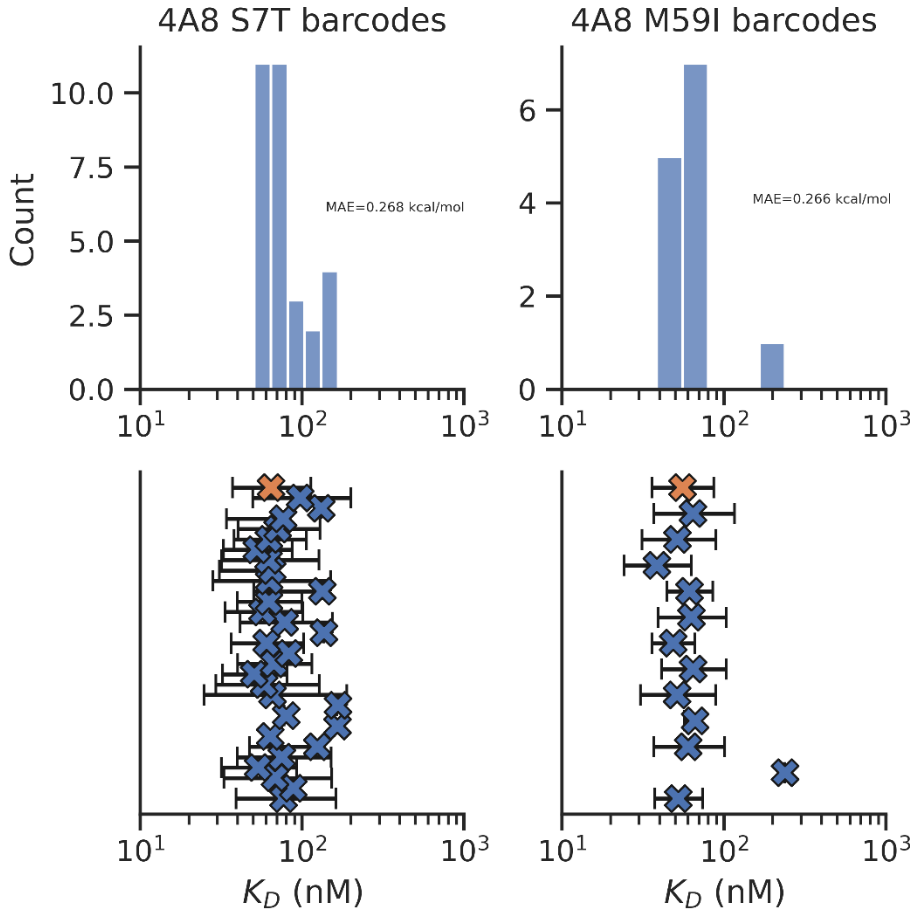

**Extended Data Figure 7 | MLE $K_D$ estimation for grouped barcodes with 95% confidence intervals.** *(Top row) Histograms of MLE $K_D$ estimates for each barcode. (Bottom row) 95% confidence intervals for each barcode (blue) and from barcodes collapsed by variant (orange)*

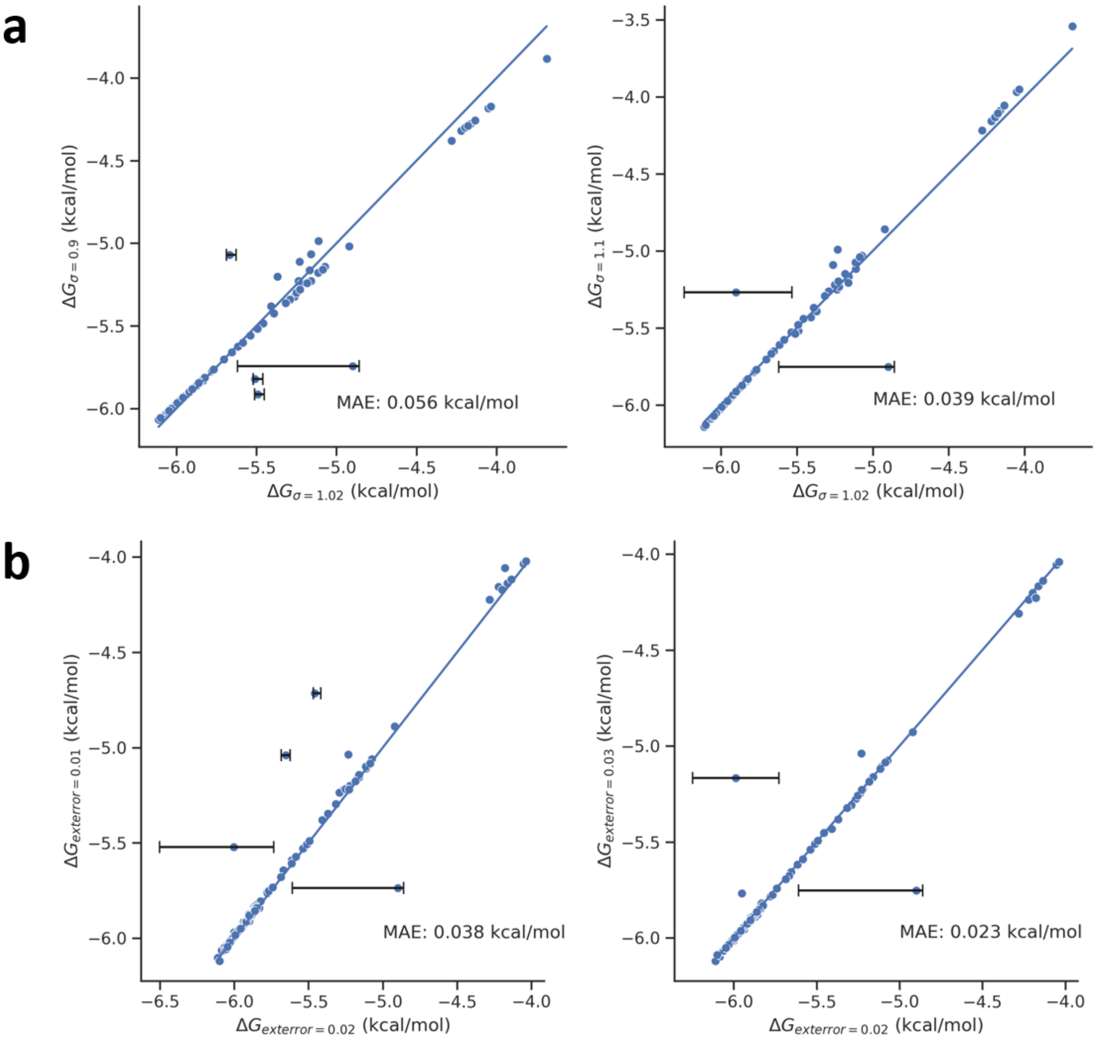

**Extended Data Figure 8 | MLE sensitivity analysis of global parameters.** *MLE calculated $\Delta G_{binding}$ values, relative to a 1 mM reference state, for 4A8 and CC12.1 data with different values of σ, the width of the isogenic lognormal distribution (Equation (2)) (a) and extrinsic error (Equation (15)) (b) with 95% confidence intervals for outliers (outliers defined as >0.3 kcal/mol MAE). Data were filtered to remove low counts and non-converged values.*

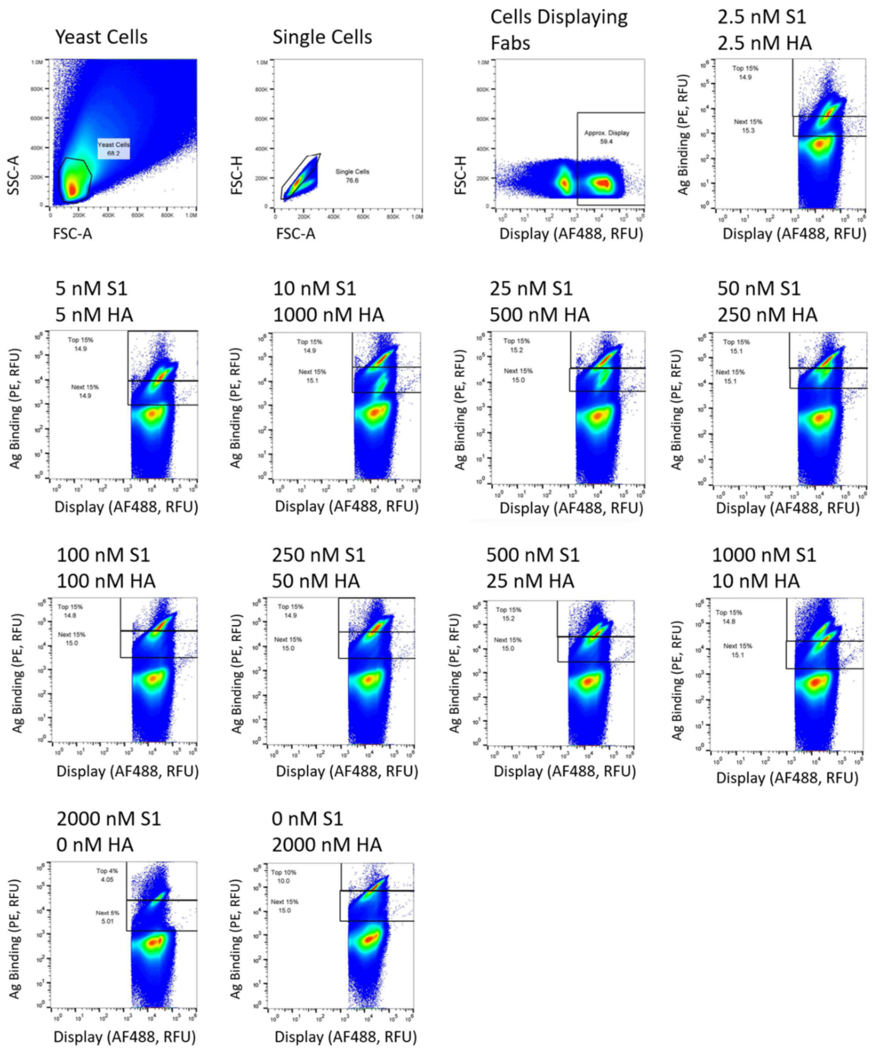

**Extended Data Figure 9 | Cytograms from YL008 mixed antigen sort with S1 and HA.** *Cytograms showing sorting gates for mixed antibody, mixed antigen sort.*

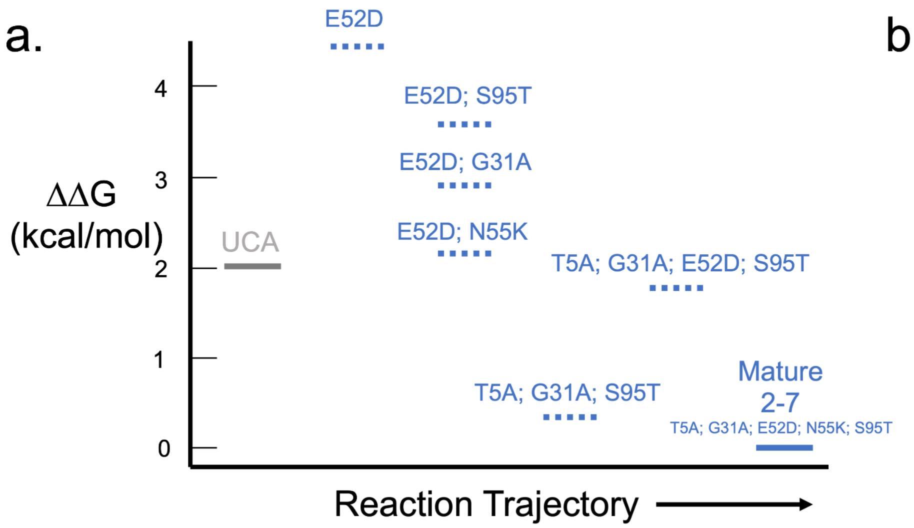

b.

| MUTATION | ΔΔG (kcal/mol) |
|---|---|
| A5T | 0 |
| A31G | 1.9 |
| D52E | -1.6 |
| K55N | 1.4 |
| T95S | 0.4 |

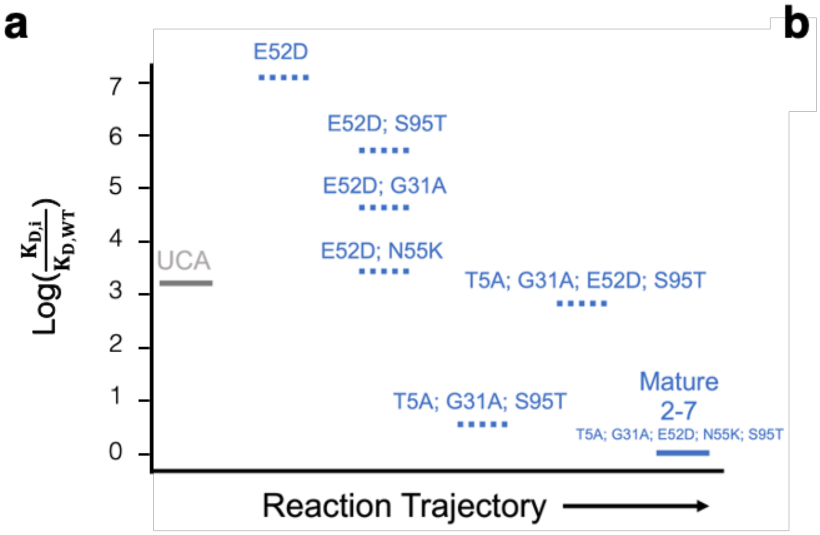

b

| Mutation | $Log(\frac{K_{D,i}}{K_{D,WT}})$ |
|---|---|
| A5T | 0.0 |
| A31G | 3.1 |
| D52E | -2.6 |
| K55N | 2.3 |
| T95S | 0.6 |

**Extended Data Figure 10 | Potential development trajectory for SARS-CoV-2 antibody 2-7.** (a) Sampling of 6 of the 30 potential intermediates between the UCA and mature 2-7. The affinity of each variant is shown as ΔΔG (kcal/mol) relative to the mature 2-7 antibody. Mature 2-7 has an inferred $K_d$ of 9.6 nM and the UCA a $K_d$ of 255 nM. (b) LASSO regression one body weights for ΔΔG for the five $V_L$ mutations. Weights in kcal/mol are shown relative to the mature 2-7 Ab. The E52D mutation is energetically unfavorable and unlikely to have appeared except in conjunction with the N55K mutation.

**References**

1. Kowalsky, C. A. *et al.* High-resolution sequence-function mapping of full-length proteins. *PLoS One* **10**, 1–23 (2015).
2. Kowalsky, C. A. *et al.* Rapid fine conformational epitope mapping using comprehensive mutagenesis and deep sequencing. *J. Biol. Chem.* **290**, 26457–26470 (2015).
3. Cossarizza, A. *et al.* Guidelines for the use of flow cytometry and cell sorting in immunological studies (second edition). *Eur. J. Immunol.* **49**, 1457–1973 (2019).
4. Boder, E. T. & Dane Wittrup, K. Optimal screening of surface-displayed polypeptide libraries. *Biotechnol. Prog.* **14**, 55–62 (1998).
5. Whitehead, T. A. *et al.* Optimization of affinity, specificity and function of designed influenza inhibitors using deep sequencing. *Nat. Biotechnol.* **30**, 543–548 (2012).
6. Klesmith, J. R., Bacik, J. P., Michalczyk, R. & Whitehead, T. A. Comprehensive Sequence-Flux Mapping of a Levoglucosan Utilization Pathway in E. coli. *ACS Synth. Biol.* **4**, 1235–1243 (2015).
7. Wittrup, K. D., Tidor, B., Hackel, B. J. & Sarkar, C. A. *Quantitative fundamentals of molecular and cellular bioengineering*. (Mit Press, 2020).
8. Johnson, M. L. Why, when, and how biochemists should use least squares. *Anal. Biochem.* **206**, 215–225 (1992).
9. Gibson, D. G. *et al.* Enzymatic assembly of DNA molecules up to several hundred kilobases. *Nat. Methods* **6**, 343–345 (2009).
10. Engler, C. & Marillonnet, S. Golden Gate Cloning - DNA Cloning and Assembly Methods. **1116**, 119–131 (2014).
11. Strawn, I. K., Steiner, P. J., Newton, M. S. & Whitehead, T. A. A method for generating user-defined circular single-stranded DNA from plasmid DNA using Golden Gate intramolecular ligation. *Biotechnol. Bioeng.* 2022.11.21.517425 (2022). doi:10.1101/2022.11.21.517425
12. Kirby, M. B., Medina-Cucurella, A. V., Baumer, Z. T. & Whitehead, T. A. Optimization of multi-site nicking mutagenesis for generation of large, user-defined combinatorial libraries. *Protein Eng. Des. Sel.* **34**, 1–10 (2021).
13. Wrenbeck, E. E. *et al.* Plasmid-based one-pot saturation mutagenesis. *Nat. Methods* **13**, 928–930 (2016).
14. Kirby, M. B. & Whitehead, T. A. Facile Assembly of Combinatorial Mutagenesis Libraries Using Nicking Mutagenesis. *Methods Mol. Biol.* **2461**, 85–109 (2022).
15. Bloom, J. D. An experimentally determined evolutionary model dramatically improves phylogenetic fit. *Mol. Biol. Evol.* **31**, 1956–1978 (2014).
16. Medina-Cucurella, A. V *et al.* User-defined single pot mutagenesis using unamplified oligo pools. *Protein Eng. Des. Sel.* **32**, 41–45 (2019).
17. Medina-Cucurella, A. V & Whitehead, T. A. Characterizing Protein-Protein Interactions Using Deep Sequencing Coupled to Yeast Surface Display. *Methods Mol. Biol.* **1764**, 101–121 (2018).
18. Haas, C. M., Francino-Urdaniz, I. M., Steiner, P. J. & Whitehead, T. A. Identification of SARS-CoV-2 S RBD escape mutants using yeast screening and deep mutational scanning. *STAR Protoc.* **2**, 100869 (2021).

19. Schirmer, M., D'Amore, R., Ijaz, U. Z., Hall, N. & Quince, C. Illumina error profiles: Resolving fine-scale variation in metagenomic sequencing data. *BMC Bioinformatics* **17**, 1–15 (2016).
